# Stochastic Decision Policies Are Optimal Under Costly Control

**DOI:** 10.64898/2026.10.06.757200

**Authors:** Jialin Li, Kenway Louie, Aadith Vittala, Marcelo G. Mattar, Paul Glimcher, Bo Shen

## Abstract

Human and animal decisions are often noisy. This variability is usually attributed to noise defects that an ideal decision maker would eliminate. Here we ask whether stochastic choice can instead be optimal once the cost of implementing prescribed policies is counted. We treat the decision maker as an ideal observer that maximizes expected gain but pays costs to sharpen its policy away from a random default. Using sequential sampling, a setting in which the optimal policy is precisely defined, we show that this trade-off makes a stochastic policy optimal. It replaces the hard decision boundaries of classical theory with soft stopping zones. This soft policy aligns with the strategy used in cutting-edge artificial learning agents for robustness. Noise that corrupts the decision-maker’s belief, an important confound of noisy behavior, produces superficially similar yet distinctive signatures in behavior. Leveraging this, we recovered the policies of two macaques with low-dimensional recurrent neural networks and found that their behavior carries both a stochastic policy and belief noise. These results establish that stochastic decision policies can themselves be optimal, offering a common principle for understanding behavioral variability in biological and artificial decision makers.

**Significance:** Humans and animals often make different choices from one to the next even facing identical evidence. This variability is usually treated as noise that an ideal decision maker would eliminate. We show that when precise control over action selections carries a cost, choosing stochastically is the optimal strategy. This turns the hard decision boundaries of classical theory into soft stopping zones. We further show that a stochastic policy and noisy evidence, although they look alike, leave distinguishable behavioral signatures, and we find both in monkey decisions. These findings establish that behavioral variability is partly a rational allocation of limited control, with implications for interpreting choice data in neuroscience, psychology, and machine learning.

## Introduction

A banal, yet fundamental observation is that biological decision-making is often stochastic. Given identical decision evidence, humans and animals choose differently from one occasion to another. This variability is usually attributed to noise: fluctuations in sensory encoding, evidence integration, or motor output that contaminate an otherwise deterministic decision process^8,18,20^. In this view, stochastic choice is a defect that an ideal decision maker would eliminate. An alternative is that stochasticity is part of the policy itself^24,36,41,62,69^. When precise control over action is costly, a decision maker may do best by choosing to act stochastically. The later view changes behavioral variability from inevitable contamination to a choice of the agent. Yet these two accounts are rarely distinguished, because both produce similarly graded behavior.

Sequential sampling offers an opportunity to separate these two accounts, taking advantage of its precisely defined optimal policy. In the past literature, decisions are commonly modeled as evidence accumulating toward a threshold - a useful framework that has unified behavioral and neural accounts of choices^6,25,51,54^. The drift diffusion model (DDM) explains variability in choices and response times through noisy accumulation to boundaries^50,51,58^. The sequential probability ratio test (SPRT) instead establishes such boundaries normatively, as stopping rules that achieve a given accuracy with the fewest samples^64,65^. With a cost per sample in a Bayesian formulation, the boundaries become the exact solution that balances the expected benefit of more evidence against the cost of obtaining it^2^. Later normative work refines this framework by showing these boundaries shifting with sampling cost and prior probability^17,28^ and collapse under time pressure^17,22^. All of these accounts share one assumption that the boundaries are hard: once accumulated evidence reaches a boundary, the agent implements the stopping rule deterministically.

Biological decision systems, however, are unlikely to meet this idealization. From a perspective of neural circuits, categorical choices emerge from competition between noisy recurrent populations. The same decision input can resolve into different winners across trials^55,66^. From a behavioral perspective, selecting a prescribed action requires cognitive control, such as maintaining task rules, suppressing prepotent responses, and resolving conflict. Control is metabolically and computationally costly. It is therefore thought to be allocated rather than applied freely ^56,57^. An agent may know the appropriate action, yet fail to implement it consistently, due to many factors such as fatigue or low motivation^7,35^.

Luckily, cost of control shares its root with information theory and can be quantified. The mental process that maps a graded belief from a random policy to prescribed categorical actions changes entropy. The entropy is maximal for a uniformly random policy and zero for a deterministic one. Sharpening a policy away from a random default removes entropy, which carries a minimal thermodynamic cost^38^. Normative theories built on this principle trade expected reward against the information cost of turning a default policy into a selective one^36,37,48^. Once the cost of control is incorporated into the current scenario of sequential sampling, a deterministic policy implemented at the hard boundaries may no longer be optimal. Instead, the optimal policy carries some degree of stochasticity, leading to probabilistic stopping near decision boundaries evident as soft-stopping zones rather than hard boundaries.

Uncovering such a stochastic policy from behavior is challenging due to the confounds of other sources of variability. Noise in sensory encoding, inference, memory, and evidence integration distorts the agent’s internal belief^8,18,21,30^. Noisy belief leads to apparently similar graded choice, even when the agent adopts a deterministic policy during evidence accumulation. Graded choice has long been described with logit or softmax rules^40,42,45,59^, but these rules are agnostic about where the variability arises. Here we distinguish two sources. First, belief noise arises upstream of the policy. It causes the internal belief to deviate from the ground-truth evidence, so that trials with identical evidence carry different internal beliefs. This dispersion of belief around the evidence arises even though the mapping from belief to action may remain deterministic. Second, stochastic policy is an explicit strategy of the agent during action selection. A certain internal belief can yield different actions across trials under a stochastic policy. Other than these two, variability arising after an action is selected, for example, noise during motor execution^20^, falls outside our scope.

Here we ask whether a stochastic policy is optimal once the cost of control is counted in the normative theory, and whether it can be dissociated from the graded behavior caused by belief noise. We derived the analysis based on a cue-based sequential-sampling task in which each cue carries a precisely defined likelihood ratio^34,70^. We formulated the task as a partially observable Markov decision process (POMDP) and derived the optimal policy under belief noise and costly control of selecting action policies.

The normative theory reveals that belief noise and stochastic policy, although superficially similar when behavior is plotted against ground-truth evidence, produce distinct temporal signatures in behavior. In particular, because belief noise enters with every sample, the dispersion of the internal belief grows over time; whereas a stochastic policy acting on an accurate belief produces no such dispersion but creates a bounded floor of stochastic behavior even at extremely certain belief. These signatures make the two sources of stochasticity identifiable from behavior alone. We test these predictions in two macaques performing the same task, using low-dimensional recurrent neural networks (tiny RNNs)^31^ to recover each animal’s decision strategy. The inferred policies reveal behavioral signatures of stochastic policy that lead to soft stopping zones instead of hard decision boundaries; belief noise coexists and exhibits its distinct feature of dispersion that accumulates over time. Together, these findings renew the normative theory of sequential decision-making by distinguishing variability in constructing a decision belief from stochasticity in selecting action policies. They recast the classical framework of deterministic threshold crossing with a resource-rational account in which stochastic stopping policies are themselves optimal.

## Results

### A cue-based sequential sampling task and its conventional optimum

We studied a cue-based sequential sampling task in which evidence arrives one cue at a time and each cue carries a known likelihood ratio between two hypotheses^34,70^ (**Fig. 1A**). Because the evidence conveyed by every cue is exactly defined, the task admits an optimal policy and allows us to compare the behavioral impacts of distinct sources of stochasticity.

**Figure 1:**
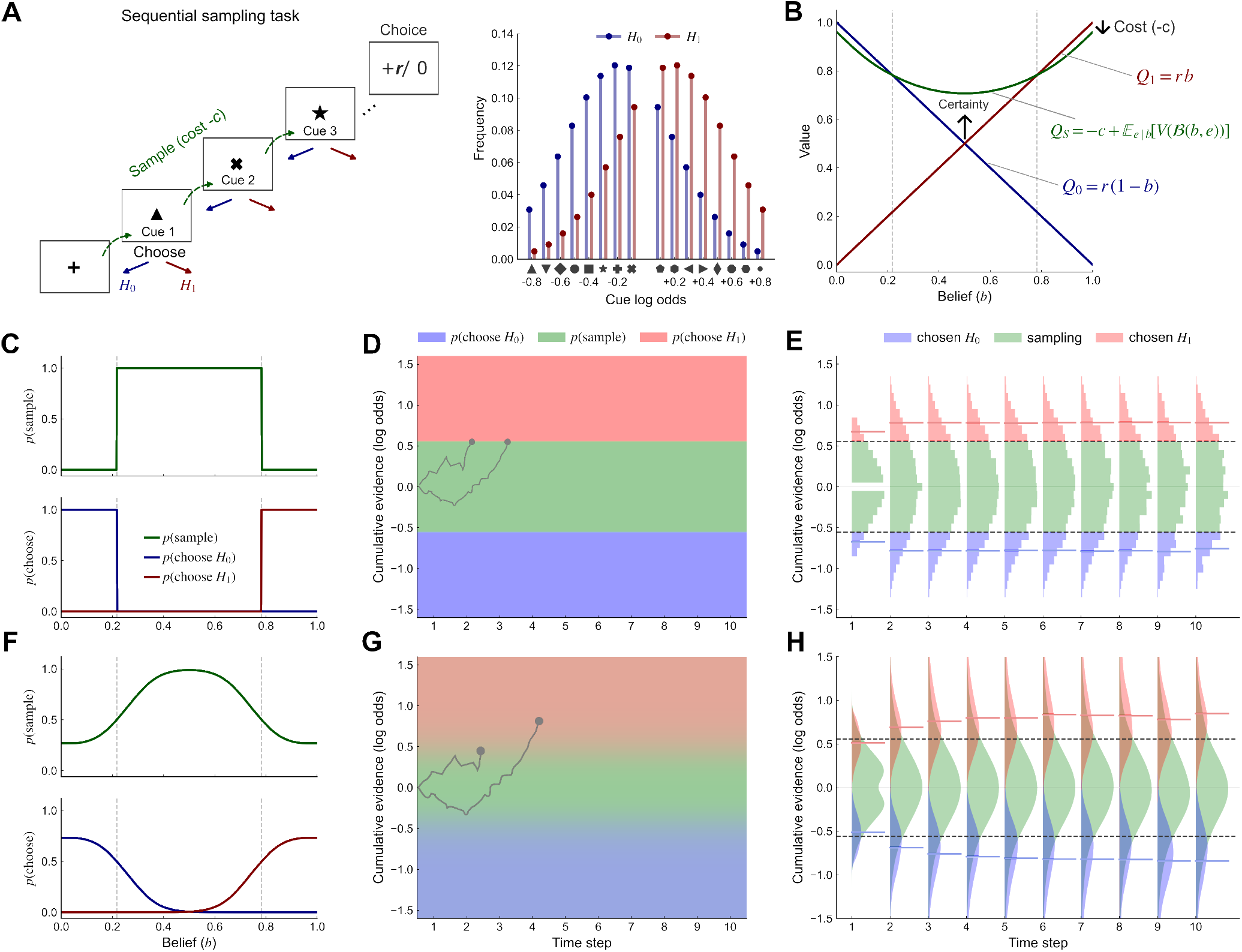
Sequential sampling task, action values, and deterministic versus stochastic policies. (**A**) Left: cue-based sequential sampling task. At each cue, the agent either samples another cue (green; cost *c* per sample) or commits to *H*_0_or *H1* (blue/red), receiving reward *r* = 1 for a correct choice and 0 otherwise. Right: cue frequencies differ between the two world states; numbers on the x-axis give the log_10_ likelihood ratio favoring *H*_1_over *H*_*0*_ for each cue. (**B**) Action values of a time-unconstrained ideal observer as a function of belief *b = P(H*_*1*_): *Q*_*1*_ (choose *H*_*1*_, red), *Q*_*0*_ (choose *H*_*0*_, blue), and *Q*_*S*_ (sample, green). Dashed vertical lines mark the optimal decision boundaries. (**C**) Deterministic policy: *p*(sample) (top) and *p*(choose *H*_*0*_), *p*(choose *H*_*1*_) (bottom) switch sharply at the boundaries (dashed). (**D**) Deterministic policy over time and cumulative evidence. Blue, green, and red regions indicate choosing *H*_*0*_, sampling, and choosing *H*_*1*_; dashed lines mark the boundaries. Visualization was shown up to *t* = 10. (**E**) Distributions of cumulative evidence on sampling (green), choose-*H*_0_ (blue), and choose-*H*_1_ (red) trials from 50,000 simulated trials are separated at the boundaries (dashed, from D). Overlaid segments mark the mean evidence at commitment. (**F-H**) As in C-E for a stochastic (soft) policy *(τ* = 0.04). Action probabilities vary continuously with belief, with dashed lines marking *p*(sample) = 0.5 (F); the policy field blends colors according to the local action probabilities (G); and the evidence distributions (kernel density estimates) of sampling and commitment trials overlap near the soft stopping zones (H).

At the start of each trial, a hidden world state *w* ∈ {0,1} is drawn with equal probability, and cues are generated from the corresponding state-specific distribution. At each time step, the agent either commits to one of two hypotheses, *H*_0_ or *H*_1_, or samples another cue at cost *c*. A correct commitment yields reward *r* and an incorrect one yields nothing. The agent must therefore decide not only which hypothesis to choose, but also whether the expected benefit of another cue justifies its cost.

We formulated the task as a partially observable Markov decision process (POMDP)^2,5,9,17,49,60,61^, in which the agent’s belief *b = P*(*H*_i_ | *e*_1:*t*_) accumulates all evidence received. The expected gain of committing immediately (**Fig. 1B**, blue and red) is

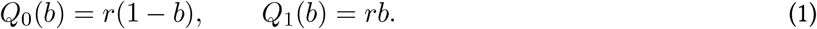

The value of sampling one more cue (green) is obtained by dynamic programming,

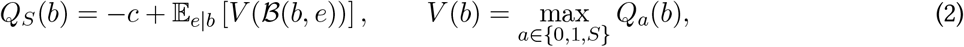

where the agent calculates the expected values for the future steps given their current belief by assuming maximizing gain at every possible situation and updates their belief as an ideal Bayes observer *(Methods)*. To illustrate such a dynamic process, an additional cue is most informative when the agent is most uncertain (*b* = 0.5) and least informative near certainty (*b* → 0 or *b →* 1), where it can no longer repay its cost. Consequently, *Q*_*S*_ exceeds both commitment values only within an intermediate range of beliefs (**Fig. 1B**).

For an ideal observer, who maps belief to action without error, the optimal action maximizes the action value *(Proofs*, Sections 1–2),

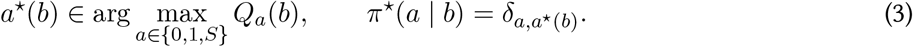

This policy is deterministic: *p*(sample) equals 1 within the region when belief is uncertain and 0 outside it, and the switch occurs exactly where *Q*_*S*_ intersects *Q*_0_ or *Q*_*1*_ (**Fig. 1C**).

Applied recursively over time, the same rule defines decision boundaries in belief or, equivalently, in cumulative log-likelihood ratio (log LR, base 10 throughout; *Methods)*. We first consider a time-unconstrained case, in which the agent may sample indefinitely, and relax this assumption later. Without a deadline, the optimal policy is stationary and its boundaries are constant over time (**Fig. 1D**). The agent samples while its belief remains in the middle region of continuation and commits as soon as the belief crosses either boundary, i.e. the threshold-crossing mechanism standard in the literature. As a result, the cumulative evidence on sampling and commitment trials occupies disjoint ranges, partitioned exactly at the boundaries (**Fig. 1E**). This hard partition is the benchmark against which we evaluate stochastic agents in the following text.

### Stochastic action selection replaces hard boundaries with soft stopping zones

The conventional optimum assumes that the agent converts belief into action without error. We relaxed this assumption by perturbing the action values,

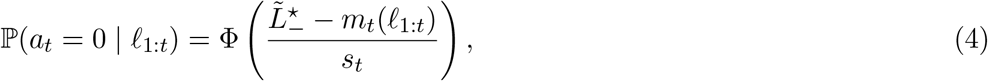

with the agent selecting the action of largest *U*_*a*_ (*b*). The perturbation *n*_*a*_ takes the form of random utility theory^40,42–44^. Different from that, here we specifically refer to this as stochasticity in action selection, where the agent maps from belief to action. We refer to the resulting policy as a stochastic, or soft, policy. It is distinct from belief noise, which corrupts the belief upstream of this mapping (see analysis in the following text).

When *η*_*a*_ are i.i.d. type-I extreme-value (Gumbel) variables, choice follows the multinomial logit^40,43^ *(Proofs*, section 3),

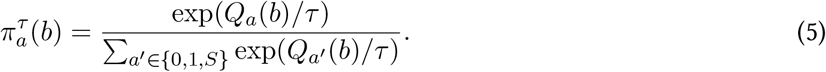

The temperature η sets the degree of stochasticity: smaller η sharpens transitions between actions, larger η spreads probability across actions, and η *→*0 recovers the deterministic policy of Eq. 3.

Under a stochastic policy, the same belief yields different actions across trials. The step of **Fig. 1C** becomes a graded transition: as belief increasingly favors either hypothesis, *p*(sample) declines smoothly while *p*(choose) rises (**Fig. 1F**). In cumulative-evidence space, the hard boundaries give way to soft stopping zones in which sampling and commitment coexist (**Fig. 1G**). Accordingly, the evidence distributions of sampling and commitment trials overlap (**Fig. 1H**). This overlap distinguishes a stochastic policy from the conventional deterministic one, and we use it below to diagnose macaque behavior. These features do not depend on the Gumbel form; they hold for general perturbations of the action values with a continuous distribution *(Proofs*, Section 3).

### A stochastic policy is optimal under a cost of control

Precise control over action is not free, since shaping behavior away from a default response requires effort^36,37,56^. We therefore asked whether a stochastic policy becomes optimal once this cost is counted. Remaining agnostic about its neural implementation, we quantified the cost of control from information-theoretic principles^36–38,48^: concentrating probability on specific actions requires control relative to an uncontrolled default.

Here we took the default to be the uniformly random policy, since it corresponds to the perturbation *n*. with infinite stochasticity,

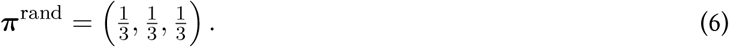

At such an extreme, the uniformly random policy has maximal entropy. A deterministic policy, as assumed by the conventional optimum, places all probability on one action and has zero entropy, i.e., the opposite extreme.

We quantified the control required to shift a policy away from the default by the Kullback-Leibler (KL) divergence,

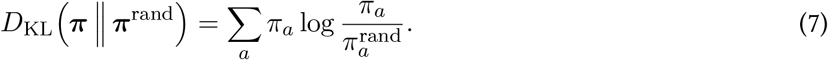

The KL form generalizes to non-uniform defaults^36^ and inherits the thermodynamic interpretation of information processing^38,48^; for the uniform default used here, it equals the entropy reduction relative to the random policy. The agent maximizes the regularized value

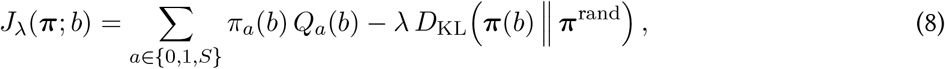

where the first term is the expected gain and the second is the cost of control. The price *λ*> 0 converts information (in nats) into units of reward and reflects the biophysical efficiency with which a system can sharpen its policy: smaller λ makes precise control cheaper, whereas larger λ makes it more expensive.

Maximizing Eq. 8 yields *(Proofs*, Section 4)

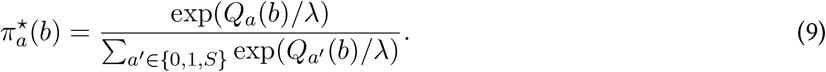

Under costly control, the optimal policy is therefore stochastic and takes exactly the logit form of Eq. 5, with temperature equal to the price of control, *τ= λ*. The logit rule is thus not an ad hoc model of choice noise but itself the optimal policy of an agent that pays for control.

**Fig. 2A** shows the geometry of this family in the probability simplex *p*_*0*_ + *p*_*1*_ + *p*_*S*_ = 1. Each curve traces the policy at a fixed τ as belief varies from *b* = 0 to *b* = 1. At low τ, the policy runs along the edges and vertices of the simplex, reflecting nearly deterministic switching among choosing *H*_*0*_, sampling, and choosing *H*_*1*_. As*τ* increases, the curves bow into the interior, and as *τ* → ∞ they collapse onto the centroid, where choice no longer depends on belief. For any finite *τ* > 0, the curves remain bounded away from the edge *p*_*S*_ = 0, even at *b* = 0 or *b* = 1, an essential signature that distinguishes from belief noise shown below.

**Figure 2:**
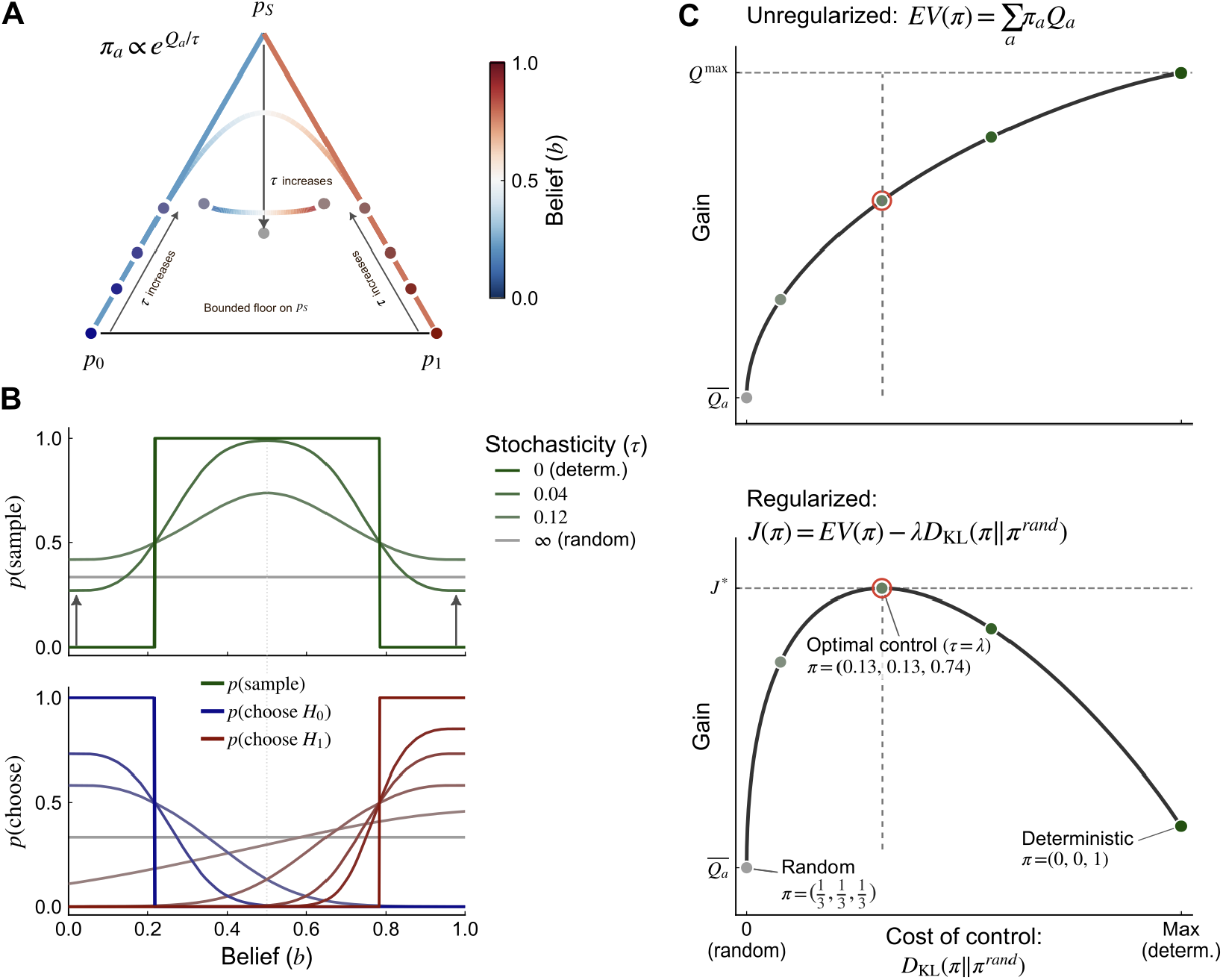
The optimal policy softens under a cost of control. (**A**) Policy simplex over the three actions (*p*_0_, choose *H*_0_; *p*_1_, choose *H*_1_; *p*_*S*_, sample). Each curve traces the logit policy π (· | *b)* as belief *b* varies (colorbar) at a fixed temperature. As *τ* increases from 0 (deterministic) toward œ (random), the curves bow away from the simplex edges toward the centroid. (**B**) *p*(sample) (top) and *p*(choose) (bottom) as functions of belief for six temperatures (graded green). Smaller τ yields steeper, more deterministic policies; larger τ flattens the policy toward the random baseline (gray). For every finite τ, *p*(sample) remains above zero at extreme beliefs (Eq. 10). (**C**) Cost of control, *D*_***KL***_*(π* || ***π***^rand^) at *b =* 0.5, increases from left to right; policy stochasticity, *H*(***n***) = log 3 − *D*_*KL*_, decreases correspondingly. Because *D*_*KL*_ decreases strictly with τ, positions along the axis correspond one-to-one to temperatures, from τ *→ ∞* (random, left) to τ →0 (deterministic, right). Top: the unregularized expected gain *EV*(*n*) rises from 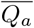 (random) toward *Q*^max^ (deterministic). Bottom: the regularized value *J_λ_* (π) is maximized by a soft policy at intermediate control, where τ =λ (red circle, dashed lines). Dots mark five example policies; three are annotated with their action probabilities τ = *(p*_*0*_, *p1,p*_*S*_).

**Fig. 2B** shows the same family as a function of belief. Increasing *τ* flattens all action probabilities, and *p*(sample) remains above zero even at extreme beliefs, in contrast to the staircase of the deterministic policy. The height of this floor follows directly from Eqs. 1-2. For example, as *b* → 1, *Q*_1_ → *r, Q*_*0*_ → 0, and *Q*_*S*_ → *r* − *c*. Hence

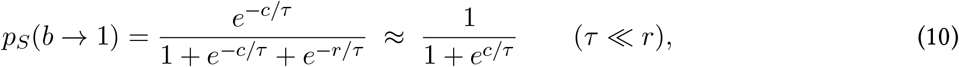

and symmetrically as *b →* 0. The sampling floor is therefore set by the ratio of sample cost to temperature. It vanishes only as *c/τ* → ∞, and for a fixed degree of stochasticity a larger sample cost makes it less apparent.

**Fig. 2C** plots the expected gain and the regularized value against the cost of control, ***D***_*KL*_(π|| ***π***^rand^) (shown at *b* = 0.5). For the uniform default, this cost equals the reduction in policy entropy, *D*_*KL*_ = log 3 − *H*(π), so moving rightward from the random policy (*τ* → ∞, left) to the deterministic policy (*τ* → 0, right) increases control and decreases policy stochasticity. The expected gain rises monotonically with control and reaches its maximum only in the deterministic limit. The regularized value instead peaks at an intermediate level of control, where *τ* = λ. Policies to the right of this peak (*τ* < *λ*) earn more reward but pay disproportionately to suppress policy entropy; policies to the left (*τ* > *λ*) save on control but forgo more reward than they save.

This result leads to a reinterpretation of stochasticity. Behavioral stochasticity need not be an imperfection of a noisy system. Once the cost of control is counted, the stochastic policy is the exact optimum, and its degree of stochasticity matches the price of control.

### Belief noise mimics a stochastic policy but leaves distinct signatures

Graded behavior does not necessarily reflect a stochastic policy. Noise in sensory encoding, evidence integration, or memory can corrupt the belief before it reaches the policy^8,18^. We modeled this belief noise as Gaussian perturbations of the internal log-odds that accumulate across samples, 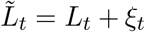 with *ξ*_*t*_~*N*(0, *σ*^2^*t*), where *L*_*t*_ is the ground-truth cumulative log LR and the internal belief is 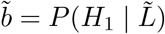 *(Methods)*. Because the variance of *ξ*_*t*_ grows linearly with *t*, trials that share the same ground-truth evidence *L*_*t*_ hold internal evidence 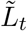 dispersed around it with standard deviation 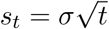. Thus, this dispersion widens with every sample.

In the agent’s own coordinates, the optimal policy under belief noise remains a hard threshold (**Fig. 3A**; *Proofs*, Section 5), and the internal evidence on sampling and commitment trials is cleanly partitioned by the boundaries (**Fig. 3C**). Experimenters, however, observe the ground-truth evidence *L*_*t*_, not 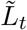. Referred to the ground-truth posterior *b*, the hard internal policy is convolved with this dispersion of internal evidence 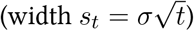 and becomes a smooth function of belief (**Fig. 3B**). The evidence distributions of sampling and commitment trials then overlap (**Fig. 3D**), as they do under a stochastic policy (**Fig. 1H**). Overlap alone therefore cannot identify the source of stochasticity.

**Figure 3:**
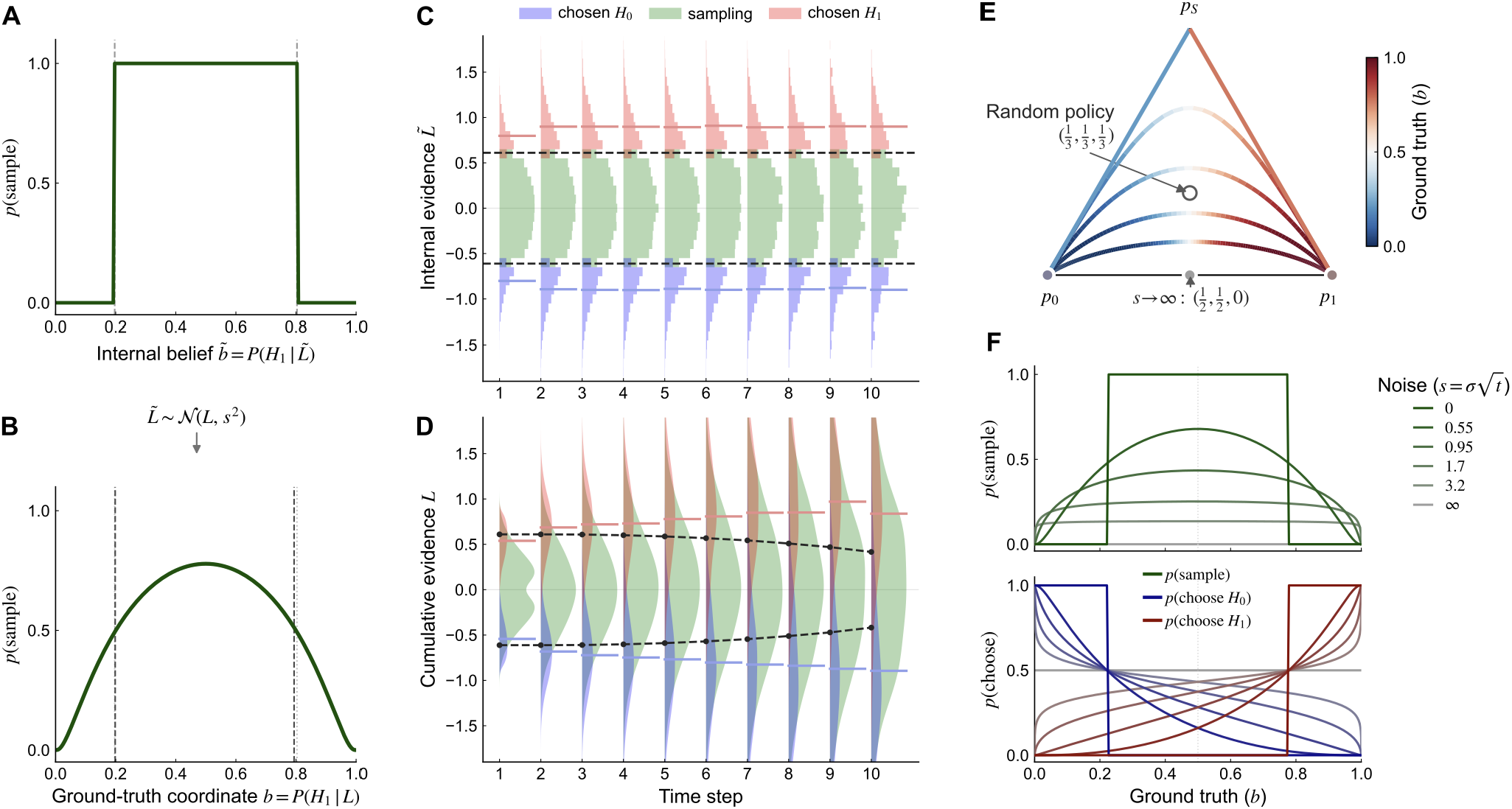
Belief noise makes a hard internal policy appear soft. (**A**) In the agent s internal belief 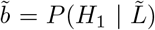, the optimal policy is a hard threshold: *p*(sample) = 1 within the continuation region and switches to 0 once the internal evidence crosses either boundary 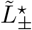(dashed). The internal evidence 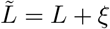 with *ξ*_*t*_~*N*(0, *σ*^2^*t*), deviates from the ground-truth evidence *L*. (**B**) Referred to the ground-truth posterior *b = P(H1* | *L*), the same hard policy is convolved with a Gaussian of width 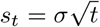, yielding a smooth *p*(sample) even though the internal policy is unchanged. (**C**) Internal view of 50,000 simulated trials. Distributions of internal evidence 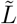 across time steps, colored by action (blue, choose *H*_0_; green, sample; red, choose *H*_1_), are sharply separated at the boundaries 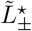 (dashed). Overlaid segments mark the mean evidence at commitment. (**D**) The same trials against the ground-truth evidence *L*. Distributions overlap across actions; dashed segments mark the per-step *p*(sample) = 0.5 crossings, which broaden over time as the dispersion *s*_*t*_ grows. (**E**) Policy curves in the simplex *(p*_*0*_, *p*1, *p*_*S*_) for increasing belief noise 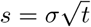, colored by belief *b*. As noise grows, the curves migrate toward the *p*_0_-*p*_i_ edge and their endpoints accumulate at 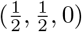 (filled marker), unlike the stochastic-policy limit at the centroid 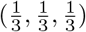 (open marker). (**F**) *p*(sample) (top) and *p*(choose) (bottom) against the ground-truth posterior *b* for increasing belief noise. As *s* grows, the peak of *p*(sample) falls toward 0 rather than settling at 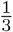 (dashed line), the stochastic-policy limit.

Three signatures nonetheless separate the two sources (**Table 1**). First, they have different limits. As belief noise grows, the policy curves in the simplex migrate toward the edge *p*_*S*_ *=* 0, with endpoints accumulating at 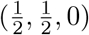 (**Fig. 3E**), whereas a stochastic policy converges on the centroid 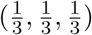 (**Fig. 2A**). Second, they differ at extreme evidence. Belief noise drives *p*(sample) to zero whenever the evidence is strong enough to make the internal belief extreme, so the peak of *p*(sample) falls toward zero as noise grows rather than a floor above zero (**Fig. 3F**); a stochastic policy instead maintains the floor of Eq. 10. Third, they differ in time. Because belief noise enters with every belief update, its dispersion accumulates over time. Thus, the ground-truth evidence distributions flatten over successive time steps (**Fig. 3D**), and the *p*(sample) curves broaden and their peaks fall (**Fig. 4C**). A stochastic policy acts only once per step at the mapping from belief to action, which makes its *p*(sample) curves stationary (**Fig. 4B**). A corollary of the third signature is that, at a fixed level of ground-truth evidence, trials differ in their internal belief under belief noise, so the trial-wise policy disperses across trials and this dispersion grows with *t*. Under a stochastic policy acting on an accurate belief, the trial-wise policy is a fixed function of the ground-truth evidence.

**Table 1:**
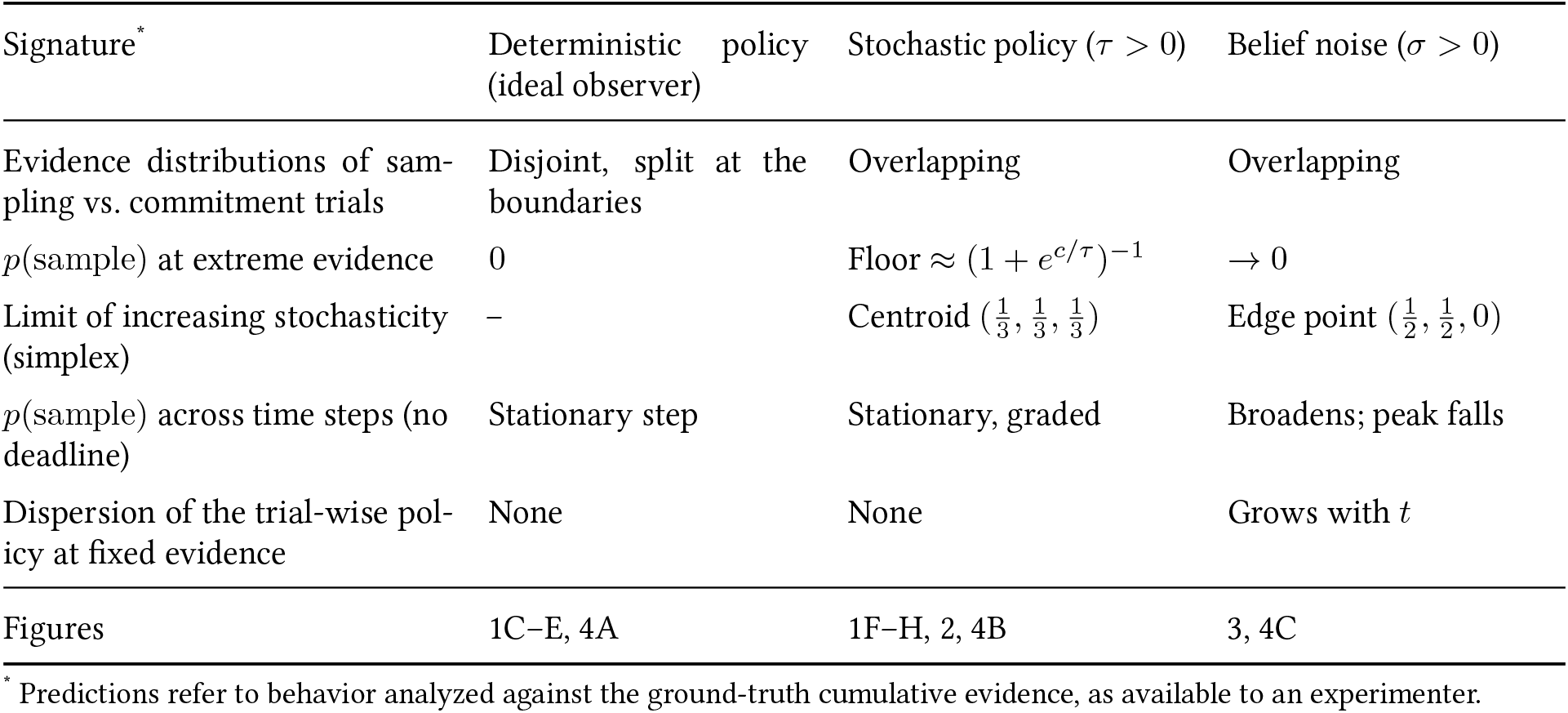
Behavioral signatures that distinguish sources of stochasticity in sequential sampling.

| Signature<sup>*</sup> | Deterministic policy (ideal observer) | Stochastic policy (*τ* > 0) | Belief noise (*σ* > 0) |
|---|---|---|---|
| Evidence distributions of sam-pling vs. commitment trials | Disjoint, split at the boundaries | Overlapping | Overlapping |
| *p*(sample) at extreme evidence | 0 | Floor ≈ (1 + *e*<sup>*c*/*τ*</sup>)<sup>−1</sup> | → 0 |
| Limit of increasing stochasticity (simplex) | – | Centroid $(\frac{1}{3}, \frac{1}{3}, \frac{1}{3})$ | Edge point $(\frac{1}{2}, \frac{1}{2}, 0)$ |
| *p*(sample) across time steps (no deadline) | Stationary step | Stationary, graded | Broadens; peak falls |
| Dispersion of the trial-wise pol-icy at fixed evidence | None | None | Grows with *t* |
| Figures | 1C–E, 4A | 1F–H, 2, 4B | 3, 4C |

<sup>\*</sup> Predictions refer to behavior analyzed against the ground-truth cumulative evidence, as available to an experimenter.

**Figure 4:**
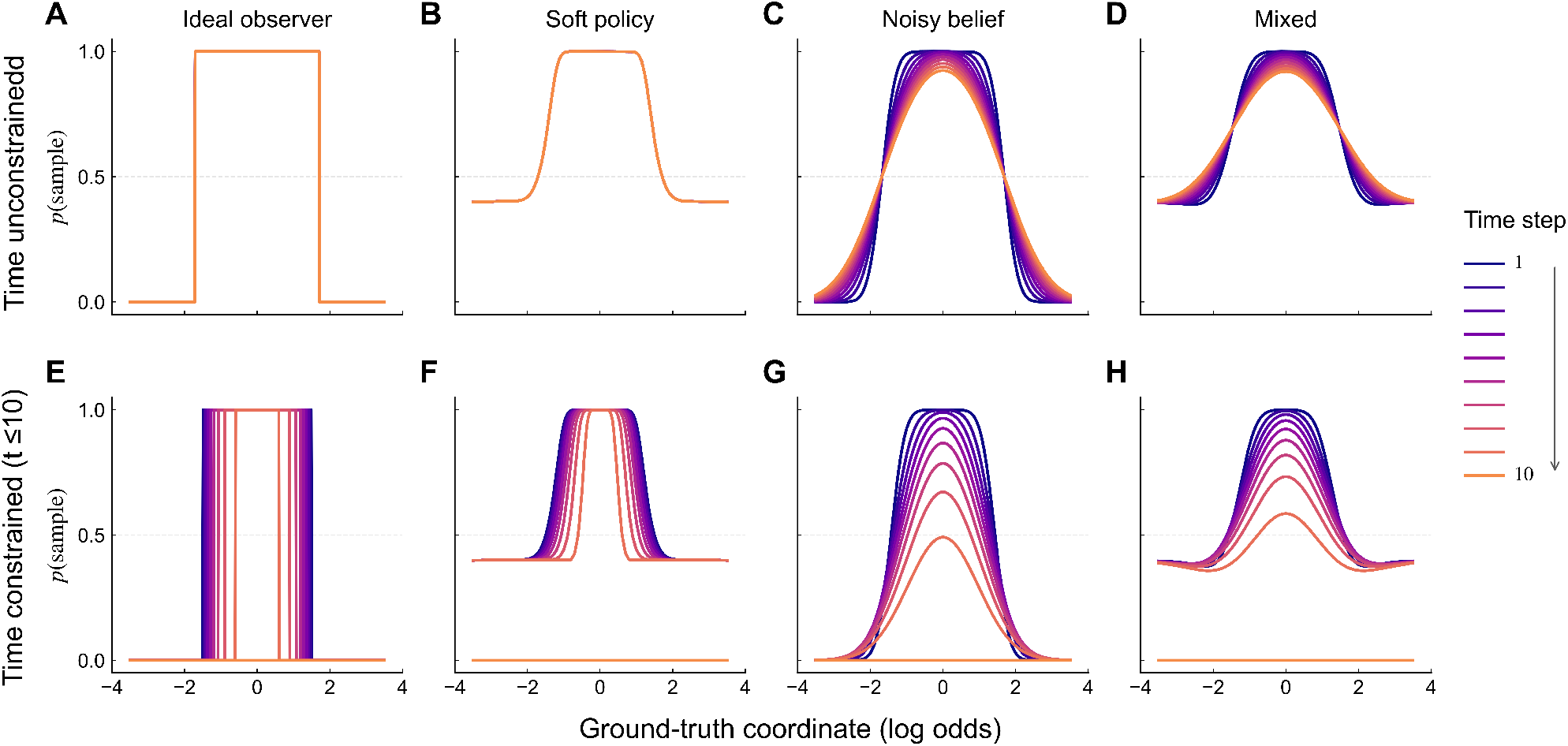
Temporal signatures of sampling policies. Each panel shows the analytical *p*(sample) as a function of ground-truth cumulative evidence (log LR) for time steps *t* = 1 to 10 (dark to light; colorbar). Gray dashed lines mark *p*(sample) = 0.5. Top row (**A-D**): time-unconstrained condition. Bottom row (**E-H**): time-constrained condition, in which no reward can be earned after *t* = 10 but each sample still incurs its cost. Columns correspond to four agents: the ideal observer (**A, E**); a stochastic policy (**B, F**; *τ* = 0.01); belief noise (**C, G**; *a* = 0.1); and both sources combined (**D, H**; *τ* = 0.01, *a* = 0.1). (**A**) The ideal observer’s policy is a step delimited by the optimal boundaries and is stationary, so curves overlap across time steps. (**B**) A stochastic policy softens the step into a graded transition and bounded above zero at extreme evidence and is stationary over time. (**C**) Belief noise blurs the policy along the ground-truth evidence axis; because the dispersion of internal evidence accumulates as 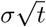, the curves broaden and their peaks fall over time. (**D**) Combining both sources yields the floor of B together with the time-dependent broadening of C. (**E-H**) The same agents under the time constraint. As the deadline approaches, the value of sampling collapses, the continuation region narrows, and *p*(sample) falls toward zero at every level of evidence by *t* = 10, favoring commitment on progressively weaker evidence. D and H show non-monotonicity on *p*(sample) near the two ends of extreme belief due to the non-monotonicity of distance between *Q*_*S*_ and *Q*_*0/1*_ when ground-truth evidence approaches the two ends. Sample cost for simulation *c* = 0.004.

Belief noise therefore produces graded behavior even when the agent applies a deterministic policy, but it leaves signatures that allow the two sources of stochasticity to be dissociated from behavior alone.

### The signatures persist over decision contexts

Humans and animals often decide under an external deadline or an internal sense of urgency^12,17^. We therefore extended our analysis to impose a deadline after which no reward can be earned (after *t* = 10), while each sample still incurs its cost *(Methods)*. **Fig. 4** compares *p*(sample) without (top) and with (bottom) the deadline. For the ideal observer, the deadline narrows the continuation region as it approaches (**Fig. 4E** versus **A**), reproducing the collapsing boundaries described in the literature^17,22^. The same collapse occurs under a stochastic policy (**Fig. 4F**) and under belief noise (**Fig. 4G**). *p*(sample) is driven toward zero as the deadline approaches, and at *t* = 10 the agent no longer samples at any level of evidence. The signatures nevertheless survive at earlier time steps: the sampling floor of the stochastic policy (**Fig. 4F**) and the progressive broadening under belief noise (**Fig. 4G**) remain visible. A deadline thus reshapes the policy without erasing the features that identify its source.

In biological decision makers, both sources of stochasticity likely coexist. An agent with both a stochastic policy and belief noise inherits the signatures of each. Without a deadline, its *p*(sample) curves are elevated by the sampling floor and broaden over time (**Fig. 4D**); with a deadline, both signatures are superimposed on the deadline-driven collapse (**Fig. 4H**).

Together, these analyses show that a stochastic policy and belief noise both produce graded behavior when analyzed against ground-truth evidence, but through different mechanisms. Belief noise makes a deterministic internal policy appear soft from the experimenter’s perspective; a stochastic policy makes the policy itself soft. Because the two leave distinct signatures, a sampling floor at extreme evidence on the one hand and time-accumulating broadening and dispersion on the other (**Table 1**), their contributions can be dissociated from choice behavior. We next applied this framework to macaque behavior.

### Tiny RNNs recover the sampling policies of macaques

We reanalyzed the behavior of two macaques (monkeys E and J) performing the same cue-based sequential sampling task^34^. The task used eight cues, with log LR values of ±0.3, ±0.5, ±0.7, and ±0.9, rather than the sixteen cues used in our simulations *(Methods)*. There was no explicit deadline, although the animals could adopt an internal deadline or urgency signal (**Fig. 4E-H**). On average, monkey E sampled 5.7 ± 1.4 effective steps before committing to a choice, whereas monkey J sampled 2.4 ± 1.5 *(Methods)*.

We first asked whether the animals’ behavior shows hard boundaries. For both monkeys, the distributions of cumulative evidence on commitment and sampling trials overlapped substantially at every time step (**Fig. 5A**). At the same level of evidence, the animals sometimes committed and sometimes continued sampling. This overlap rules out a deterministic policy acting on an accurate belief, which would predict disjoint distributions (**Fig. 1E**). It is, however, consistent with either a stochastic policy (**Fig. 1H**) or with belief noise (**Fig. 3D**), or with both. Distinguishing among these requires the policy itself, the probability of each action given the cue history, which choices alone do not reveal.

**Figure 5:**
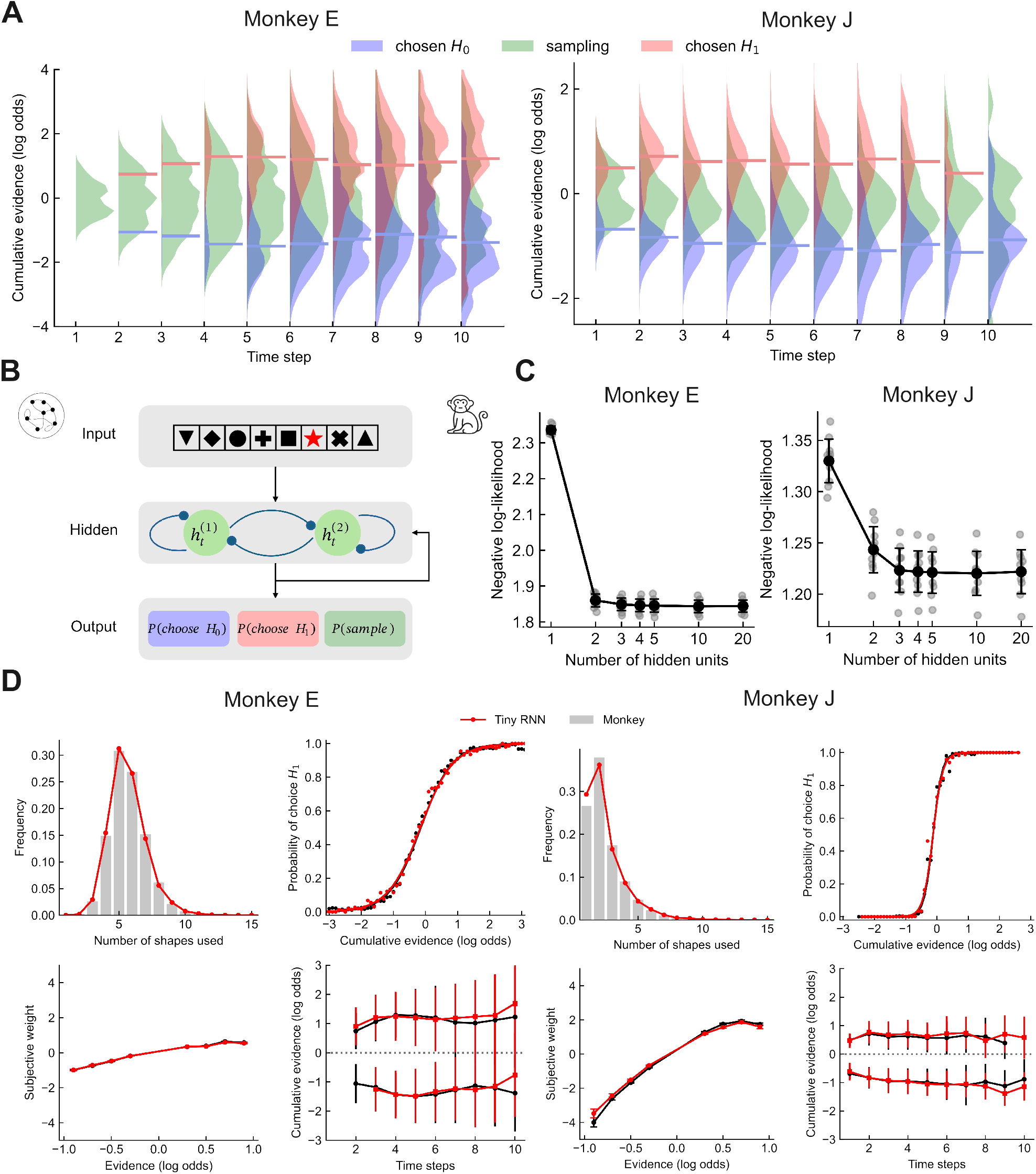
Tiny RNNs capture the sequential sampling behavior of macaques. (**A**) Distributions of cumulative evidence across time steps on choose-*H*_0_ (blue), choose-*H*_1_ (red), and sampling (green) trials for monkey E (left) and monkey J (right). Overlaid segments mark the mean evidence at commitment. In both animals, sampling and commitment trials overlap at every time step. (**B**) Tiny-RNN architecture with two hidden units. The network receives the cue sequence, updates a low-dimensional hidden state, and maps it to probabilities of choosing *H*_0_, choosing *H*_*1*_, or sampling. (**C**) Held-out trials negative log-likelihood of networks with 1, 2, 3, 4, 5, 10 and 20 hidden units, fitted and evaluated separately to monkey E (left) and monkey J (right). (**D**) Monkey behavior (gray/black) and two-unit RNN simulations (red) for monkey E (top) and monkey J (bottom). From left to right: distribution of the number of shapes sampled before commitment; psychometric function, *p*(choose *H*_1_) against cumulative evidence; subjective weight of each cue against its log odd; and mean cumulative evidence at commitment across time steps.

To recover each animal’s policy, we fit tiny recurrent neural networks (tiny RNNs)^31^ to its trial-by-trial behavior. Tiny RNNs learn a compact nonlinear mapping from cue sequences to actions without committing in advance to a particular model of the decision process, while their low dimensionality keeps the learned dynamics interpretable. Each network was a gated recurrent unit (GRU) network^10^ that received the current cue as input, updated a low-dimensional hidden state, and output a probability distribution over choosing *H*_*0*_, choosing *H*_1_, and sampling at every step (**Fig. 5B**). Networks were trained separately for each monkey by maximum likelihood (i.e. cross entropy) with Adam^33^, jointly predicting the choice and the time step of commitment *(Methods)*. Importantly, the output policy was not constrained to be graded: a network fit to deterministic behavior can saturate its action probabilities toward 0 or 1.

To avoid overfitting and measure the dimensionality of macaque’s behavior, we implemented nested crossvalidation that separates trials for training, validation and evaluation *(Methods)*. For both monkeys, increasing the hidden state from one to two units substantially reduced the held-out negative log-likelihood (monkey E: *t*(18) = 68.31, *p =* 3.39*e*^−23^; monkey J: *t*(18) = 8.3976, *p =* 1.22*e*^−07^), whereas further hidden units did not yield further improvement *(h* = 2 v.s *h* = 3: monkey E: *i*(18) = 1.32, *p* = 0.203; monkey J: *i*(18) = 1.94, *p* = 0.068, **Fig. 5C**). We therefore analyzed the two-unit networks as the most parsimonious adequate models. When simulated in the task environment experienced by the animals, the two-unit networks reproduced the behavioral features reported by Kira et al.^34^: the distribution of the number of cues sampled, the psychometric function, the subjective weight of each cue, and the mean cumulative evidence at commitment (**Fig. 5D**). One-unit networks failed to reproduce these features (**Fig. S1**), indicating that at least two latent dimensions are required to account for the animals’ behavior.

### Macaque policies carry signatures of both a stochastic policy and belief noise

We next compared the policies recovered by the tiny RNNs with the signatures predicted by the normative theory (**Table 1**).

#### A graded policy with a sampling floor

For both monkeys, *p*(sample) as a function of ground-truth cumulative evidence was bell-shaped rather than step-like (**Fig. 6A**): sampling was most likely near zero evidence and declined gradually as evidence accumulated toward either hypothesis, forming soft stopping zones rather than hard boundaries. In monkey E, *p*(sample) remained well above zero even at the largest cumulative evidence the animal encountered. This floor is the signature of a stochastic policy (Eq. 10; **Figs. 2B** and **4B**) and cannot arise from belief noise alone (**Fig. 3F**).

**Figure 6:**
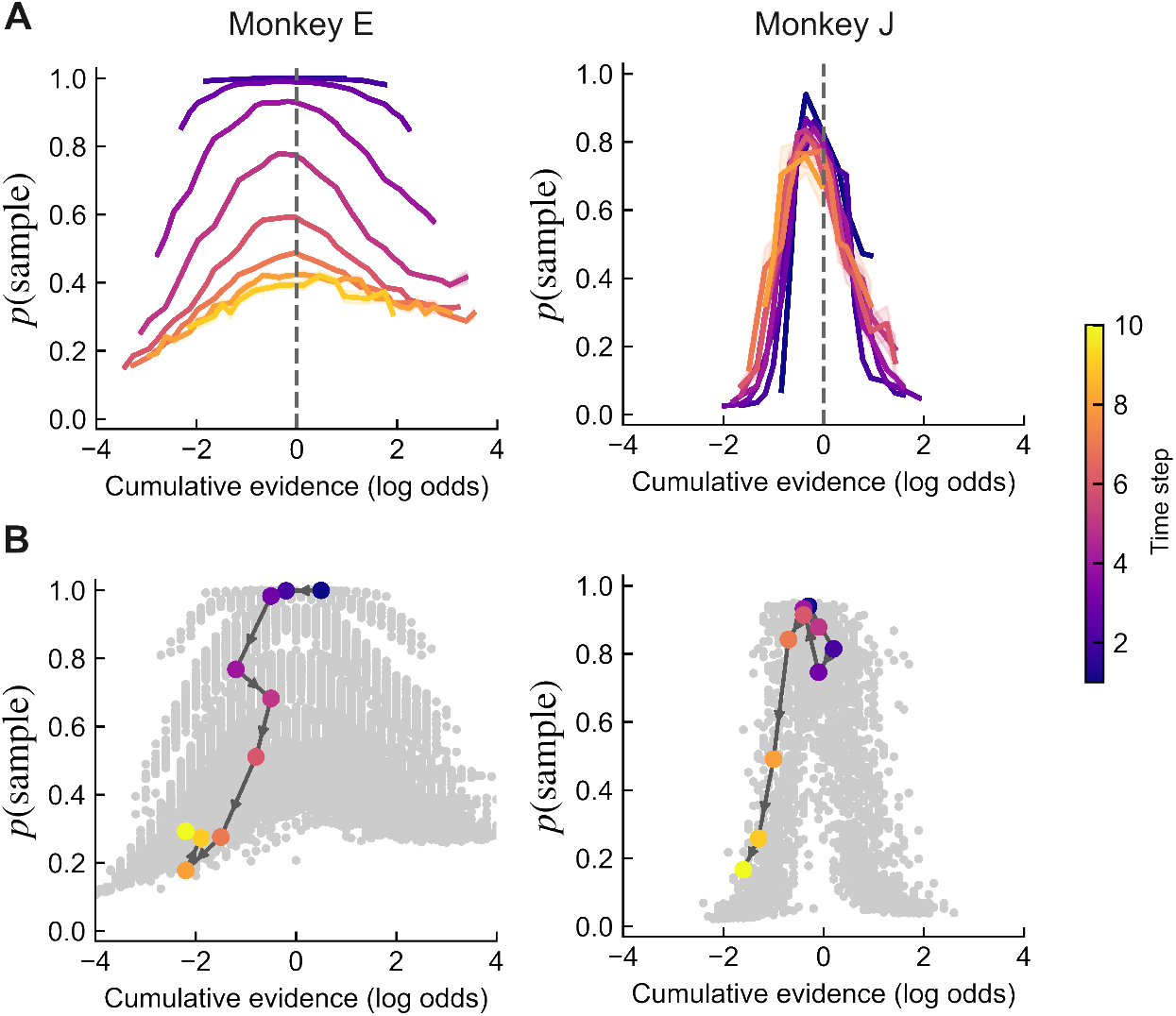
Sampling policies recovered by tiny RNNs show soft stopping zones. (**A**) mean *p*(sample) generated by the two-unit networks fit to monkey E (left) and monkey J (right) as a function of cumulative evidence at each time step. (**B**) Trial-wise *p*(sample) plotted against cumulative evidence (gray points). Colored circles trace an example trial, with color denoting time step.

Monkey J’s continuation region was narrower and its floor less apparent. This is expected from the theory. Monkey J committed after fewer samples than monkey E, consistent with a higher effective sample cost, and a larger *c*/*τ* both narrows the continuation region and lowers the floor for the same degree of stochasticity (Eq. 10).

#### Graded policies on single trials and in the networks’ internal coordinates

Trial-wise policies support the same conclusion. A deterministic policy outputs *p*(sample) of 0 or 1 at every step. Instead, the networks’ trialwise *p*(sample) spanned the full interval between 0 and 1 (**Fig. 6B**, gray points), and individual trials traversed intermediate values across successive steps (colored markers). Because the networks were free to saturate their outputs, these graded values indicate that stochasticity was required by the data rather than imposed by the architecture.

The networks’ hidden states show where this gradation arises. In both animals, hidden states evolved along a smooth, curved manifold ordered by cumulative evidence (**Fig. 7A**). Because the policy layer is a linear projection of the hidden state followed by a softmax *(Methods)*, the logit of each action is a linear function of the two hidden units, and the policy is fully determined by where the hidden state lies in the two-dimensional state space (**Fig. 7C**). The readout thus organizes the manifold into a sampling region near its center, where cumulative evidence is weak and *p*(sample) is highest, and two commitment regions toward its ends, where evidence favors *H*_*0*_ or *H*_1_. These regions are not sharply partitioned: the policy changes gradually from one region to the next, so that hidden states moving along the manifold pass through mixtures of sampling and commitment rather than switching abruptly between them. This gradation is informative about the source of stochasticity. Under belief noise alone, the theory predicts a hard threshold in the agent’s own coordinates (**Fig. 3A,C**), with softness appearing only when the policy is referred to ground-truth evidence. The hidden state is the network’s internal belief, yet the policy remains graded when expressed in these coordinates. The softness is therefore not merely inherited from a deviation of belief from evidence but resides in the mapping from belief to action itself, as expected of a stochastic policy.

**Figure 7:**
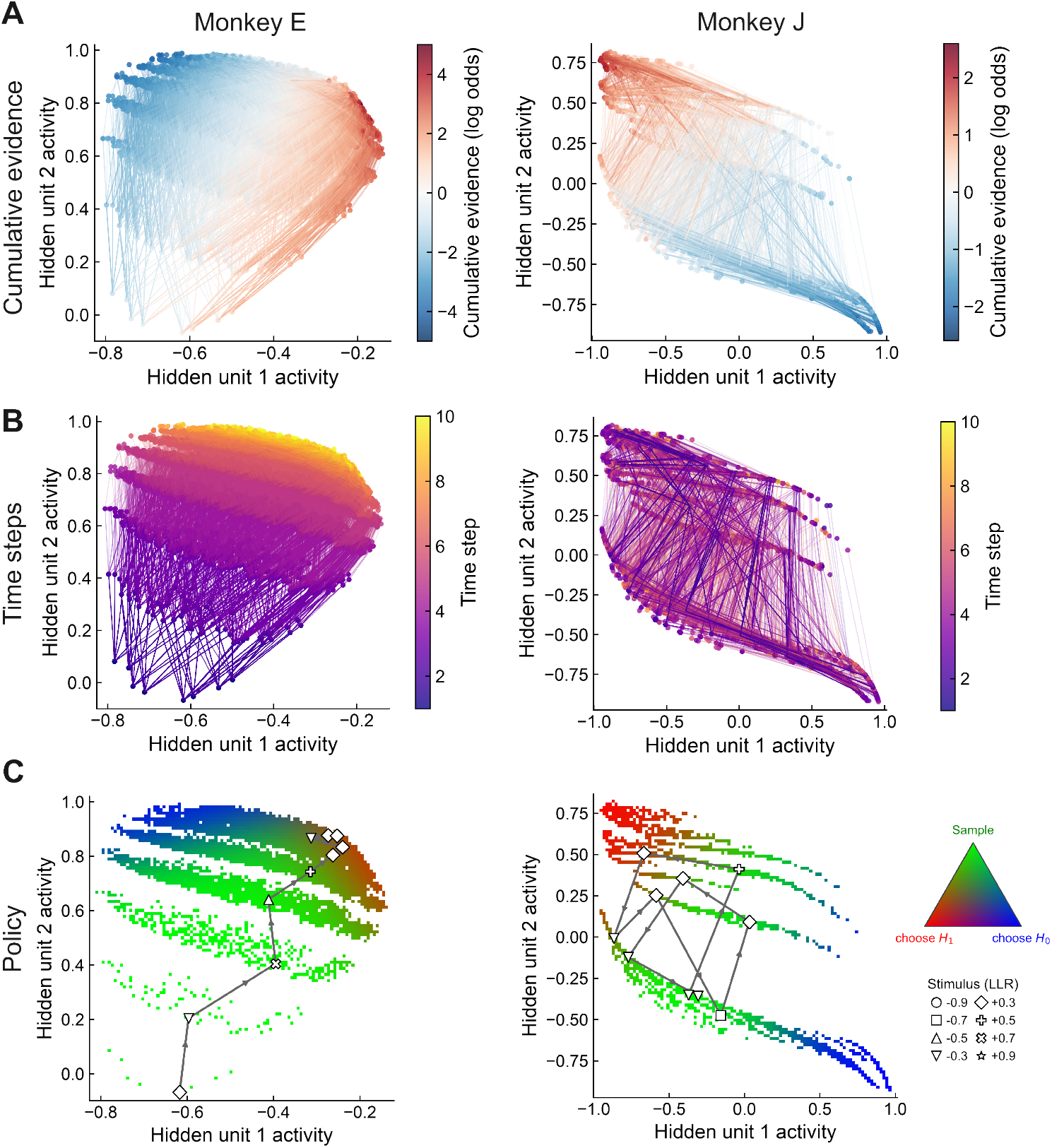
Low-dimensional hidden-state dynamics of tiny RNNs. Hidden-state trajectories of the two-unit networks fit to monkey E (left) and monkey J (right). Each point is the hidden state at one time step of a trial; lines connect successive states within a trial. (**A**) Hidden states colored by cumulative evidence (log LR). (**B**) Hidden states colored by time steps. (**C**) Hidden states colored by the network’s policy, shown as a mixture of the probabilities of sampling, choosing *H*_*0*_, and choosing *H*_1_ (ternary color key).

#### A stochastic policy under urgency and under stationarity

The temporal structure of the recovered policies differed between animals, as the theory allows. In monkey E, the *p*(sample) curves shifted downward across time steps, lowering the probability of sampling at every level of evidence and progressively favoring commitment (**Fig. 6A**, left). This is the pattern predicted under a time constraint (**Fig. 4F,H**), indicating an internal urgency signal even though the task imposed no deadline. Throughout this shift, *p*(sample) never collapsed to zero, as predicted for a stochastic policy (**Fig. 4F**). In monkey J, the curves largely overlapped across time steps (**Fig. 6A**, right), indicating a stationary policy governed primarily by cumulative evidence, as predicted for the time-unconstrained case (**Fig. 4B,D**).

The hidden-state geometry mirrors this difference. In monkey E, hidden states were separated by time step in addition to cumulative evidence (**Fig. 7B**, left), providing the representation of elapsed time needed for a timedependent policy. Accordingly, *p*(sample) in the state space was higher for states visited at earlier time steps and declined as trials progressed (**Fig. 7C**, left), the geometric counterpart of urgency. This decline remained gradual, so that urgency shifted the graded policy toward commitment rather than imposing a hard collapsing boundary.

In monkey J, hidden states from different time steps overlapped extensively, with no clear temporal stratification comparable to that observed in monkey E (**Fig. 7B**). Within this representation, the policy transitioned from sampling-dominated regions to regions favoring either choice as evidence became more extreme (**Fig. 7C**, right). This pattern is consistent with a its time-invariant and stationary policy despite substantial variation in the underlying hidden states. Under both urgency and stationarity, the policy expressed in the networks’ own coordinates remained graded - the signature of a stochastic policy.

#### Dispersion that accumulates over time

Belief noise predicts that, at a fixed level of ground-truth evidence, the trial-wise policy varies across trials and that this dispersion grows with each sample; a stochastic policy acting on an accurate belief predicts no such dispersion (**Table 1**). In both monkeys, the variability of trial-wise *p*(sample) among trials with matched cumulative evidence increased across time steps (**Fig. 8A**; Monkey E: *β*_*t*_ = 7.39*e*^−3^, *R*^*2*^ = 0.639, *p* < 1*e*^−3^; Monkey J: *β*_*t*_ = 3.28*e*^−3^, *R*^2^ = 0.536, *p* < 1*e*^−3^), and the variability of the hidden states increased likewise (**Fig. S2**). This accumulating dispersion is the signature of belief noise. The dependence of this variability on evidence further supports its origin upstream of the policy (**Fig. 8B**). Variability was low near zero cumulative evidence and at extreme evidence, and largest at intermediate evidence, within the soft stopping zones. This is the pattern expected when a dispersed belief is passed through the policy. A given spread of internal belief changes *p*(sample) little where the policy is flat, either deep in the sampling region near *L* = 0, where nearly all trials continue, or far beyond the stopping zones, where nearly all trials commit. The same spread changes *p*(sample) the most where the policy is steep, in the transition between sampling and commitment. Trial-to-trial variability of the policy at matched evidence thus scales with both the dispersion of belief, which grows with time step, and the local slope of the policy, which depends on evidence. Variability arising independently of belief would show no such dependence on evidence, and a stochastic policy acting on an accurate belief would produce no variability at all. In the fitted networks, where we did not include any noise perturbation, the networks capture the deviation of the internal state from ground-truth evidence by leveraging systematic misweighting of cues, evident in subjective weights that depart from the true log LR (**Fig. 5D**). Like random noise, such misweighting accumulates with each sample. From the experimenter’s perspective, random noise and distorted accumulation of evidence both constitute belief noise, because both corrupt the belief upstream of the policy *(Proofs*, Section 5). In the biological brain, however, both random noise and misweighting may exist.

**Figure 8:**
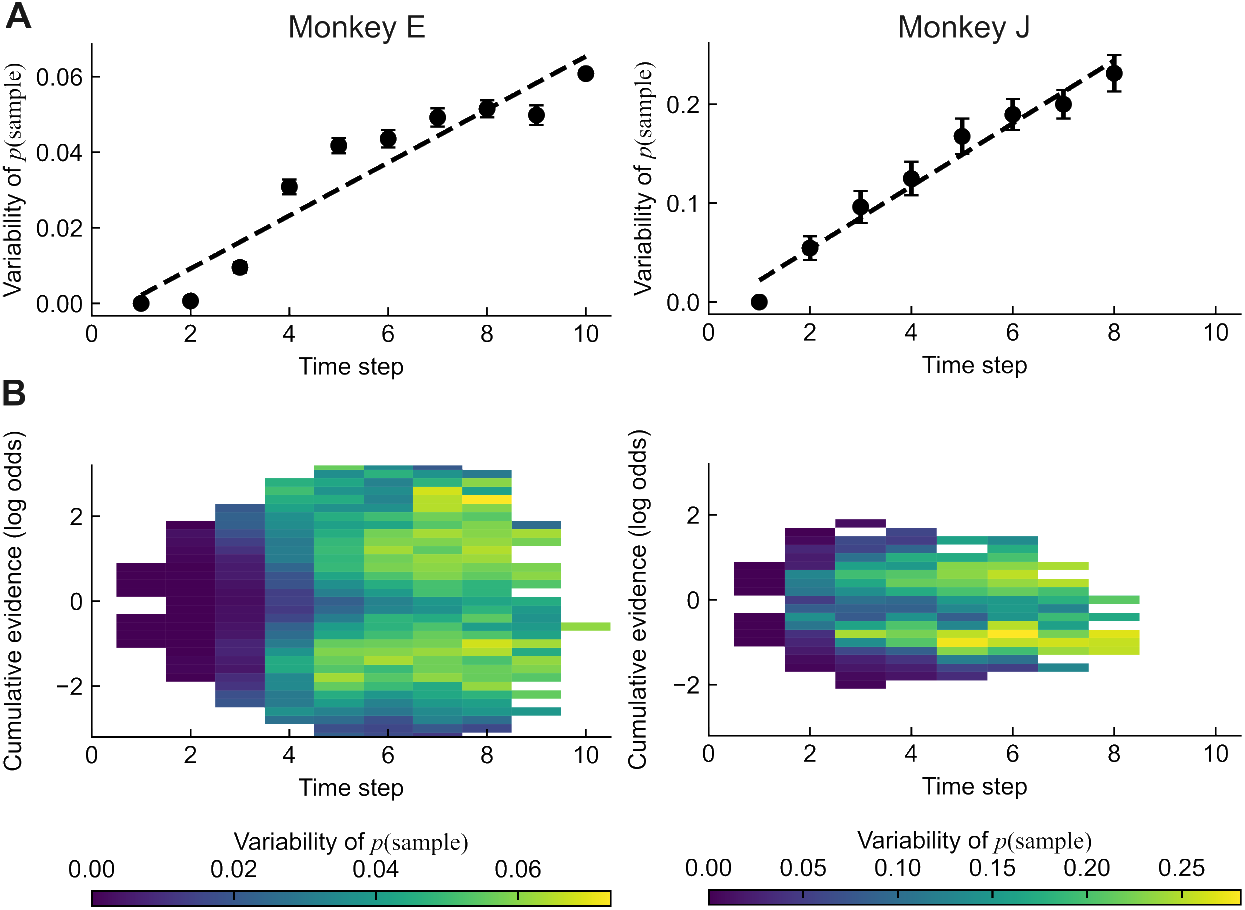
Belief noise in macaque policies accumulates over time. Variability of the trial-wise *p*(sample) among trials with matched cumulative evidence for monkey E (left) and monkey J (right). (**A**) Variability as a function of time step. In both animals, variability increases over time, as predicted for belief noise that accumulates with each sample (**Fig. 4C**; **Table 1**). Dashed line shows the fitted linear regression (Monkey E: β_*t*_ = *7*.*39e*^−3^, *p* < *1e* ^−3^; Monkey J: β_*t*_ = 3.28*e*^−2^, *p* < 1*e*^−3^). (**B**) Variability in bins of cumulative evidence over time steps. Variability is low near zero and at extreme evidence and largest at intermediate evidence, where the policy transitions between sampling and commitment.

Together, the recovered policies of both macaques show soft stopping zones rather than hard boundaries and carry the signatures of two coexisting sources of stochasticity predicted by the normative theory. A stochastic policy is revealed by the sampling floor, by graded trial-wise policies, and by a policy that remains graded in the networks’ own coordinates under both urgency and stationarity. Belief noise is revealed by dispersion that accumulates over successive samples.

## Discussion

Classical accounts of sequential sampling treat the decision boundary as hard: once the accumulated evidence reaches a threshold, the agent stops and commits^2,6,17,64^. Variability in choices and response times is then explained by noise in the accumulated evidence, which blurs an otherwise deterministic stopping rule^50,51^. Our results offer a different view. Once the control needed to sharpen a policy is counted, the optimal stopping rule is no longer a threshold but a soft stopping zone, in which sampling and commitment coexist at the same belief and their probabilities change gradually with evidence. The softmax form of this policy is not imposed as a model of choice noise; it follows exactly from the trade-off between expected reward and the cost of control, with the temperature of softmax equal to the price of control. In this sense, the logit rule that has long described graded choice^40,42,43^ acquires a normative interpretation at a specific stage of the decision process, i.e., the mapping from belief to action.

Why would an agent tolerate a soft policy when a hard one earns more reward? The first answer is given by the analysis itself: control is costly, and sharpening a policy to determinism is never free^38,48,56,57^. Near the boundaries, where the values of sampling and committing are close, a deterministic choice buys little extra reward at a disproportionate price, so relaxing control there saves resources at little cost (Fig. 2C). The second answer concerns environments richer than the task studied here. A stochastic policy is the canonical mechanism of exploration in reinforcement learning, preventing premature convergence onto exploitation^59^, and humans and animals deploy random exploration when their knowledge of the environment is incomplete^13,67^. In competitive settings, unpredictability is valuable in its own right: the equilibrium of a zero-sum game is a mixed strategy that an opponent cannot exploit^23,46,63^, and monkeys approach such mixed strategies in competitive games^4,15,39^. Rats can even switch strategically into a stochastic choice mode, under noradrenergic control of anterior cingulate cortex, when they cannot outpredict a competitor^62^. In these settings, behavioral variability is not a passive consequence of imperfect sensing or reasoning but is generated and regulated by the agent^24^. Machine learning makes the same principle explicit. Maximum-entropy reinforcement learning rewards an agent for keeping its policy stochastic, which smooths the optimization landscape, discourages premature convergence to deterministic policies, and improves robustness^1,19,26,27,71^. The optimum of this entropy-regularized objective is again a softmax over action values^26,53,71^, identical to our solution for a uniform default policy and to the multinomial logit that arises from Gumbel perturbations of utility^42^. This convergence is not a coincidence; it reflects a shared principle of trading reward against the information needed to concentrate a policy.

Behavioral variability has typically been treated holistically, as a single noise term hovering around a hard decision rule. Our analysis shows that variability arising at different stages of the decision process leaves distinguishable signatures, even when its aggregate effect on choice looks the same. Belief noise and a stochastic policy both cause sampling and commitment trials to overlap in evidence, but they diverge at extreme evidence, in their limits in the policy simplex, and over time (Table 1). In the macaque data, the dependence of policy variability on evidence provided an additional diagnostic: variability was concentrated where the policy is steep and was low where it is flat, as expected when a dispersed belief is passed through a graded policy (Fig. 8B). Both forms of stochasticity are also distinct from variability in motor execution, which can be injected by dedicated circuits and removed without disrupting the underlying motor program^20,32,47^. Previous work has partitioned the contributions of different sources of variability through experimental manipulation and modeling^18,21^. The signatures derived here provide a complementary route that operates on choice sequences alone and can therefore be applied to many existing datasets. Separating these sources matters because they have different underlying neural substrates. Belief noise reflects how evidence is encoded and integrated, whereas policy stochasticity reflects how strongly the agent controls its action selection. Individual differences that look alike in aggregate behavior, such as those between monkeys E and J, may therefore reflect different settings of these two processes.

The dissociation rests on a distinction between two views of the same behavior: the agent’s internal belief and the ground-truth evidence available to an experimenter. A policy that is hard in the agent’s coordinates becomes soft when referred to ground-truth evidence whenever the internal belief deviates from that evidence (Fig. 3A,B). Psychometric and chronometric functions measured against ground truth therefore cannot, on their own, reveal whether the agent’s policy is deterministic. The fitted networks offered access to both views. Against ground-truth evidence, their policies showed the dispersion that accumulates over samples, as the signature of belief noise. In their own hidden-state coordinates, the policy remained unnecessarily graded, as the signature of a stochastic policy (Fig. 7C). The deviation of internal belief from ground truth in the networks arose without any injected noise, through systematic misweighting of cues (Fig. 5D).

Our considerations raise a caution for machine learning. Networks trained with cross-entropy or entropy-regularized objectives typically retain some softness in their output policies, and this softness is often read as uncertainty or as a residual training error. Our analysis suggests that graded outputs in a trained network can arise from two different sources: a genuinely stochastic mapping from internal state to action, or a deviation of the internal state from the true input statistics passed through a sharper mapping. The two can be distinguished by examining the policy in the network’s own state coordinates, as we did for the tiny RNNs, and by testing whether output variability at matched inputs grows with the length of the input sequence. Conversely, the entropy coefficient used to train artificial agents provides a controlled proxy for the cost of control, which is difficult to quantify in the brain but well defined during network training. Training networks under graded entropy regularization offers a way to test directly how limited control shapes sequential decision policies.

Our theory specifies when a stochastic policy is worth adopting. The optimal policy assigns probabilities according to value differences scaled by the price of control. Softness is therefore concentrated where the values of competing actions change smoothly and cross gradually, as the values of sampling and committing do in sequential sampling (Fig. 1B). Where the reward structure changes abruptly, or where errors are catastrophic, value differences are large relative to the price of control and the optimal policy approaches determinism. Stochastic policies are thus expected to be optimal when reward changes smoothly with belief and to give way to near-deterministic control when the stakes of a single action are high. This prediction can be tested by manipulating the steepness of the reward landscape. It also connects to the neural circuits of action selection. Categorical choices emerge from competition among recurrently connected populations, in which stochastic firing allows the same input to resolve into different winners across trials^14,55,66,68^. Downstream, the basal ganglia select among competing actions^11,52^, and the sharpness of this selection is neuromodulated. Tonic dopamine has been proposed to set the effective temperature of a softmax policy^16,29^, and noradrenergic gain modulation shifts behavior between focused and labile modes^3,62^. Stochasticity in action selection may therefore be a controllable circuit parameter rather than a fixed noise floor.

### Limitations

Our framework rests on several simplifying assumptions. First, we assumed a uniform default policy. Non-uniform defaults^36^ would bias the policy without removing its graded form. Second, we modeled belief noise as accumulating Gaussian perturbations. The noise-free networks reproduced the same signature through cue misweighting (**Fig. 5D**), and distinguishing the two forms will require analysis on neural measurements^34^. Third, the cost of control was applied to the policy at each belief state, whereas action values came from the unregularized Bellman equation. Propagating this cost through a soft Bellman equation^26,71^ may shift the soft stopping zones, with effects on our signatures yet to be established. Fourth, our macaque results rest on two animals and on model-based policy estimates that needs confirmation in neural recordings. Fifth, we treated the price of control *λ* as fixed. If control is allocated by its expected value^56,57^, *λ* may vary with motivation, fatigue, or hunger^7,35^. This yields a testable prediction: larger rewards or hunger should sharpen the policy and lower the sampling floor (Eq. 10), whereas satiety or fatigue should soften it.

## Methods

### Task model and ideal observer

We formulated the cue-based sequential sampling task as a partially observable Markov decision process (POMDP). On each trial, a hidden world state *w* ∈ {0,1}, corresponding to hypotheses *H*_0_ and *H*_1_, was drawn with equal probability. At each time step *t*, the observer received a discrete cue *e*_*t*_ drawn from the state-specific distribution *f*_0_(*e*) or *f*_1_*(e)*. Each cue carries the log likelihood ratio *l*(*e*) = log_10_[*f*_1_(*e*)*/f*_0_(*e*)]. Throughout the paper, log LR and cumulative evidence are expressed in base-10 units, following the convention of the macaque studies^34,70^. The observer maintains the belief

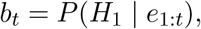

or equivalently the cumulative log-odds

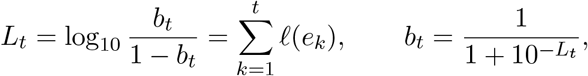

starting from *L*_0_ = 0 (*b*_0_ = 0.5). The available actions were *a* ∈ {0,1, *S*}: choosing *H*_0_, choosing *H*_1_, or sampling one more cue at cost *c*. A correct choice yielded reward *r* = 1 and an incorrect choice yielded 0.

### Cue set used in simulations

Simulations used sixteen cues with *l* ∈ {±0.1, ±0.2, &, ±0.8}. Their stateconditional frequencies were derived from an equal-variance Gaussian generative model, in which a latent cue value *x* is drawn from *N*(+*µ, s*^*2*^) under *H*_1_ and from *N(-µ, s*^*2*^) under *H*_0_, with *µ* = 2.2 and *s* = 5. Because the variances are equal, the log LR is linear in *x, l* = 2*px/*(*s*^2^ In 10). Each cue was assigned the value *x(l)* = f*s*^2^ In 10*/*(2*µ*), and *f*_*1*_*(l)* and *f*_*0*_*(l)* were set proportional to the Gaussian densities at *x(l)* under each hypothesis and normalized over the sixteen cues (**Fig. 1A**, right).

### Bellman equation

For an ideal Bayesian observer with perfect evidence representation and integration, beliefs

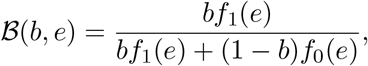

equivalently *L*_*t*_ = *L*_*t*-1_ + *l*(*e*_*t*_). The values of committing are

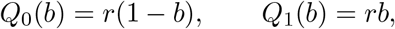

and the value of sampling is

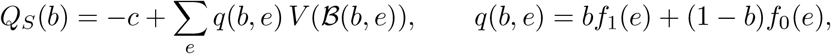

where *q(b, e)* is the predictive probability of observing cue *e* under belief *b*. The Bellman optimality equation is

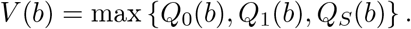

The optimal stopping boundaries 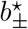 are the beliefs at which the maximizing action switches between sampling and committing to *H*_0_ or *H*_1_. In log-odds coordinates, they correspond to

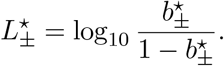

### Time-unconstrained condition

Without a deadline, we solved the Bellman equation by value iteration. We initialized *V* with the value of committing immediately,

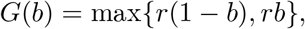

and repeatedly applied the Bellman operator

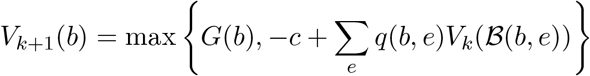

until the maximal absolute change fell below 10^−9^. This produced a stationary value function and stationary boundaries, and the same policy was applied at every time step.

### Time-constrained condition

To model a deadline after which no reward can be earned, we solved the problem by backward induction. We set

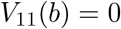

for all beliefs, so that an agent that had not committed by *t =* 10 earned nothing while still paying for each sample, and recursively computed

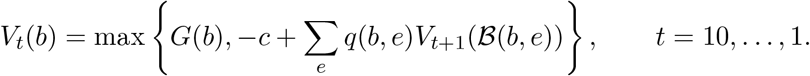

This yielded time-dependent values *Q*_*s*_,_*t*_*(b)* and boundaries 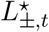 that collapse as the deadline approaches.

### Numerical implementation

Value functions were represented on a grid of 2,001 (arbitrarily picked, fine enough) equally spaced log-odds values in [—4, 4] and evaluated between grid points by linear interpolation, with values beyond the grid held at the boundary values. The boundaries were taken as the outermost grid points of the region in which *Q*_*s*_ exceeded *G*.

### Stochastic policy and cost of control

For the stochastic policy, the observer computed the same action values as the ideal observer, but selected actions according to the multinomial logit (Eq. 5),

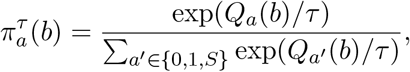

which follows from adding i.i.d. Gumbel perturbations to the action values (Eq. 4; *Proofs*, Section 3). In the time-constrained condition, *Q*_*s*_ was replaced by *Q*_*s*_,_*t*_. The stochasticity acts only at the mapping from belief to action and leaves the action values unchanged. Consequently, the points at which *p*(sample) = 0.5 lie close to the deterministic boundaries, where *Q*_*s*_ crosses *Q*_*0*_ or *Q*_1_, and coincide with them when the probability of the third action is negligible.

The cost of control was quantified as the KL divergence between the policy and the uniformly random default (Eqs. 6-7). Maximizing the regularized value *J*_*λ*_ (Eq. 8) at each belief under the constraint ∑_*a*_ *π*_*a*_ = 1 yields the softmax policy with *τ = λ* (Eq. 9; *Proofs*, Section 4). For **Fig. 2C**, we evaluated, at *b =* 0.5, the expected gain 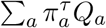, the cost *D*_*KL*_*(π*^*T*^ ||*π*^rand^) = log 3 — *H(π*^*T*^), and the regularized value *J*_*λ*_ *(π*^*T*^) across a range of temperatures *τ* for a fixed price *λ*. The policy-simplex curves (**Fig. 2A**) trace 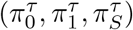 as *b* varies from 0 to 1 at fixed *τ*.

### Belief noise

For the observer with belief noise, the internal log-odds 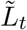 deviates from the ground-truth *L*_*t*_ through Gaussian perturbations added at every update,

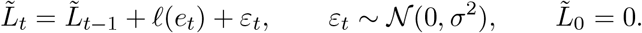

The accumulated deviation 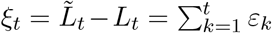 is therefore distributed as *N* (0, *σ*^*2*^*t)*, with standard deviation 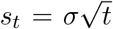. This is the belief-noise model of the Results, and *s*_*t*_ measures the dispersion of internal evidence around the ground truth. The internal belief is

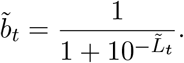

Dynamic programming was performed in the observer’s internal belief space. The values of committing are 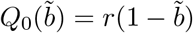 and 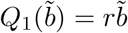, whereas the value of sampling integrates over both the next cue and the next perturbation:

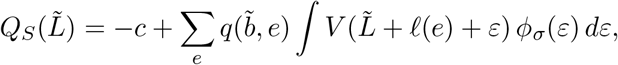

where ϕ_*σ*_ is the Gaussian density with standard deviation *σ*. The integral was evaluated by Gauss-Hermite quadrature with 401 (arbitrarily picked, fine enough) nodes. The resulting optimal policy is a hard threshold on 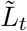, with internal boundaries 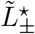 (**Fig. 3A**; *Proofs*, Section 5).

### Experimenter’s view

An experimenter observes *L*_*t*_ rather than 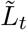. Marginalizing the hard internal policy over 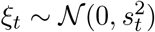 gives the policy as a function of ground-truth evidence,

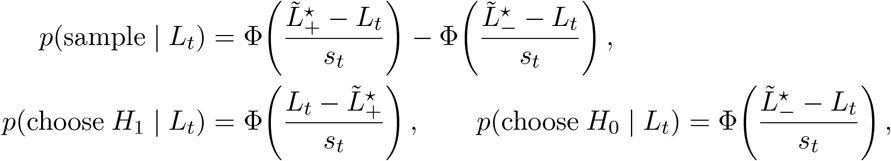

where Φ is the standard normal cumulative distribution function. In the time-constrained condition, the boundaries 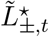 were time dependent. These analytical curves (**Figs. 3B,E,F** and **4C,G**) treat the deviation at step *t* as independent of whether a trial has survived to step *t*; the Monte Carlo simulations described below, however, do not have to make such an approximation. For the simplex and noise sweeps (**Fig. 3E,F**), the internal boundaries were held fixed while the dispersion *s* was varied.

### Combined stochastic policy and belief noise

For the observer with both sources, dynamic programming was performed in internal belief space as for belief noise, and the resulting internal action values 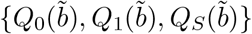 were passed through the softmax,

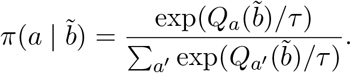

The policy as a function of ground-truth evidence was obtained by convolving this internal policy with the Gaussian distribution of *ξ*_*t*_ (width *s*_*t*_), evaluated by Gauss-Hermite quadrature. Thus, belief noise softens the policy only when it is viewed against ground-truth evidence, whereas a stochastic policy softens it also in the observer’s own coordinates.

### Simulations

For each observer, we simulated 50,000 trials of up to 10 time steps. On each trial, the world state was drawn with equal probability, cues were drawn from the corresponding distribution, and both the ground-truth evidence *L*_*t*_ and, for observers with belief noise, the internal evidence 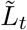 were recorded. At each step, the action was selected from the observer’s policy evaluated at its internal evidence, and the trial ended at commitment. Evidence distributions of sampling and commitment trials at each time step (**Figs. 1E,H** and **3C,D**) were computed as histograms smoothed with a Gaussian kernel. The parameters used for each figure are given in the figure legends.

### Fitting macaque behavior with tiny RNNs

#### Macaque behavioral task and dataset

We reanalyzed behavioral data from the sequential inference task reported in Kira et al. ^34^; a complete description can be found in the original study. Two rhesus monkeys (monkey E and monkey J) made binary decisions based on sequences of visual shapes. Each shape was drawn from a set of eight and was associated with a log LR of ±0.3, ±0.5, ±0.7, or ±0.9 (base 10), providing graded evidence for one of two targets. On each trial, the monkey fixated a central point while two peripheral choice targets were displayed. After a random delay, shapes were presented sequentially every 250 ms until the monkey committed to a choice by a saccadic eye movement, which it could make at any time after the first shape appeared. A correct choice was rewarded with juice if gaze remained on the chosen target for 0.2 s. There was no explicit deadline.

The dataset comprised 27,201 trials for monkey E and 17,994 trials for monkey J. Following the original study, we excluded shapes presented too close to the saccade to influence the choice, based on estimated non-decision times of 270 ms for monkey E and 180 ms for monkey J. We defined *N*_*_ as the number of effective shapes on each trial and based all analyses on the sequence from the first through the *N*_*_th shape. After this adjustment, monkey E used 5.7 ± 1.4 effective shapes and monkey J used 2.4 ± 1.5 (mean ± SD). Accuracy was 85% for monkey E and 80% for monkey J. The cumulative evidence on step *t* of a trial was the sum of the log LRs of the first *t* shapes. In the analyses, sampling at step *t* refers to steps *t* < *N*_*_, and commitment refers to step *N*_*_.

#### Network architecture

To investigate the strategy of macaques, we fitted tiny recurrent neural networks built from gated recurrent units (GRUs)^10,31^. At each time step, the current shape was encoded as a one-hot vector ***x***_*t*_ over the eight shapes, and the hidden state ***h***_*t*_ was updated as

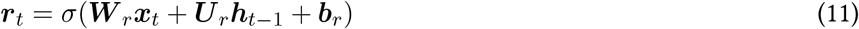

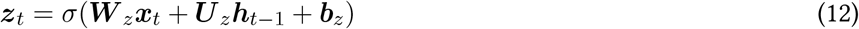

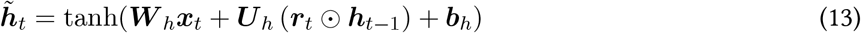

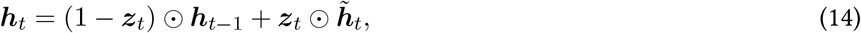

where *σ (·)* denotes the softmax function. The hidden state was projected by a linear policy layer onto three action logits, which were passed through a softmax to define the policy *π*_*θ*_ *(a*_*t*_ | ***h***_*t*_) over choosing *H*_*0*_, choosing *H*_*1*_, and sampling. The network contained no noise: given a shape sequence, its hidden states and policy were deterministic. Because the policy is defined at every step, the network jointly predicts the choice and the number of shapes sampled.

#### Maximum likelihood estimation

The networks were fit by maximum likelihood. For a trial with decision time *T* = *N*^*\**^ and observed choice *a*_*T*_ ∈ {*H*_*0*_, *H*_*1*_}, the negative log-likelihood is

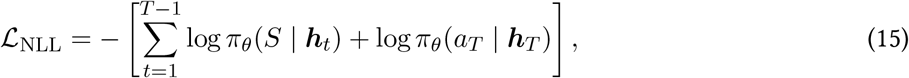

which requires the network to assign probability to sampling before the decision time and to the observed choice at the decision time. To discourage redundant connections, we added an *L*_1_ penalty on the recurrent weights ***U*** = {***U***_*r*_, ***U***_*z*_, ***U***_*h*_},

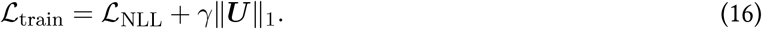

Nothing in this objective constrains the policy to be graded; a network fit to deterministic behavior can drive its action probabilities toward 0 or 1. Networks were optimized with Adam^33^ with a learning rate of 3 × 10^−3^ and a batch size of 128. The network was trained with a maximum of 2,000 epochs.

#### Nested cross validation

We evaluated model performance using nested cross validation. Specifically, trials were divided into 10 outer folds, stratified by the monkey’s final choice, to preserve the choice distribution across folds. In each fold, we use one fold for testing, while the remaining nine folds were further partitioned into 9 inner folds. Each inner model was trained on 8 inner folds and validated on the remaining fold.

For each network size *h* ∈ {1, *2*, 3, 4, 5,10, 20}, we evaluated every combination of recurrent *L*_*1*_ regularization *γ*= {10^−3^,10^−4^,10^−5^} and three network initialization seeds. For each inner fit, the checkpoint with the lowest validation loss across training epochs was retained. The validation loss was computed as the negative logprobability of the observed actions, averaged over all valid time steps. The regularization term and initialization seed were selected by the unweighted mean of this validation loss across the 9 inner folds. The selected hyperparameter configuration was then evaluated on the held-out testing fold. The predictive performance was quantified using the average trial negative log-likelihood. During training, we implemented a early stopping rule if the model has no further improvement on the validation loss after 100 training epochs, in order to prevent overfitting on the training data.

#### Estimating the dimensionality of behavior

For both monkeys E and J, RNN predictive performance improved with increasing numbers of hidden units before reaching a plateau. We operationally defined behavioral dimensionality as the smallest number of hidden units beyond which further increases in model capacity yielded no statistically significant improvement in predictive performance, as assessed using *i*-tests across outer crossvalidation folds^31^.

### Behavioral analyses of macaques and networks

#### Behavioral simulation

To test whether the fitted networks reproduce the animals’ behavior, we let the network perform 10,000 trials in an environment with the same statistics as the original experiment^34^, sampling shapes from the task distributions and actions from the network’s learned policy at each step. Behavioral analyses were performed using the two hidden unit RNN with the lowest held-out test negative log-likelihood for each monkey. All network weights were frozen throughout the simulation.

#### Psychometric curves

The psychometric curves were fitted with a logistic sigmoid for RNN’s simulated choices:

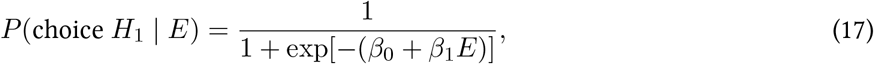

where *E* is cumulative evidence at the time of decision. The intercept *β*_0_ and slope *β*_1_ were estimated using a binomial generalized linear model.

#### Subjective weights

To estimate the subjective weight of each shape, we fit the logistic regression

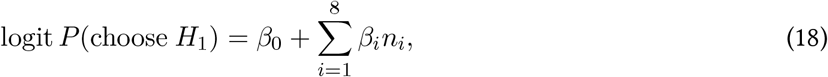

where *n*_*i*_ is the number of times shape *i* appeared among the effective shapes of a trial. The coefficients *β*_*i*_, which quantify the contribution of each shape to the choice, were compared with the ground-truth log LR of each shape, separately for the monkeys and for the networks.

#### Cumulative evidence

To smooth the cumulative-evidence distributions arising from discrete stimulus evidence, we convolved the histograms for continued sampling and each choice with a normalized five-point binomial kernel, [1, 4, 6, 4,1]*/*16, separately at each timestep (**Fig. 5A**). This kernel provides a discrete approximation to Gaussian smoothing.

#### Recovered policies analyses of the networks

To recover the policy, we used the same two-unit network as above, simulated trials with the cue sequences and recorded, at every effective step, the hidden state ***h***_*t*_ and the policy *π*_*θ*_ (· | ***h***_*t*_) from the output layer. The trial-wise *p*(sample) is the network’s sampling probability at a given step of a given trial (**Fig. 6B**). The mean policy at each time step (**Fig. 6A**) was obtained by binning trials by cumulative evidence and averaging the trial-wise *p*(sample) within each bin.

To quantify the dispersion of the policy, we grouped steps with matched cumulative evidence, that is, steps falling in the same evidence bin at the same time step, and computed the variability of the trial-wise *p*(sample) within each group (at least 10 observations):

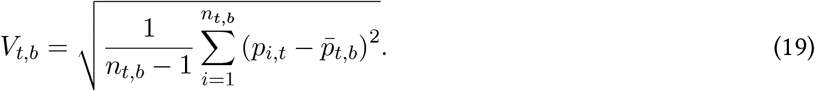

where *t* is the timestep, *b* is the cumulative evidence, *i* is the trial index within the group. Variability *V*_*t*_,_*b*_ was then summarized as a function of time step by averaging across evidence bins (**Fig. 8A**) and as a function of cumulative evidence (**Fig. 8B**). We then fitted a linear regression model for each monkey:

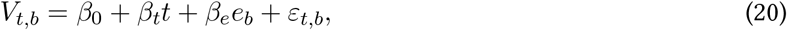

here *e*_*b*_ denotes the signed cumulative-evidence value at the bin center. Each valid bin contributed one equally weighted observation. Model fit was assessed using *R*^*2*^ and the *F*-test.

#### Hidden state analyses

For the hidden-state analyses (**Fig. 7**), each hidden state activation from these simulated trials was projected to two dimensional space colored by the cumulative evidence (A), the time step (B), or the policy (C). In C, the color was a mixture of the colors for choosing *H*_*0*_ (blue), choosing *H*_*1*_ (red), and sampling (green), weighted by the corresponding action probabilities.

The variability of hidden states among steps with matched cumulative evidence was computed in the same way, which was defined as the square root of the summed sample variances across hidden units (**Fig. S2**):

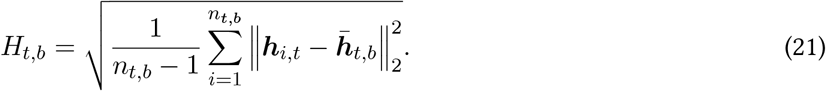

Here, ***h***_*i,t*_ is trial *i*’s hidden-state vector, 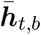 is the group’s mean hidden state, and *n*_*t*_,_*b*_ is its number of observations. For a two-unit network, this is

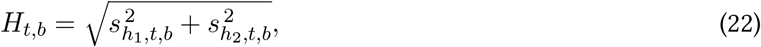

the square root of the sum of the two units’ sample variances. Larger values indicate that different stimulus histories produce more dispersed hidden representations despite matching timestep and cumulative evidence. Under a policy that depends only on ground-truth evidence, this variability is zero; it becomes nonzero when the network’s internal state, and hence its policy, differs among trials with the same evidence. Again, an ordinary least-squares linear regression model was then fitted to each monkey:

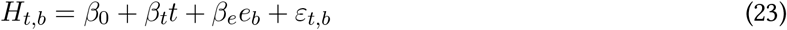

## Acknowledgments

We thank Dr. Shinichiro Kira and colleagues for sharing the macaque data. This work was supported by National Natural Science Foundation of China (32671436) to B.S.

## Author Contribution

Conceptualization: B.S., Methodology: J.L., M.M., and B.S., Code: J.L. and B.S., Normative analysis: J.L., A.V., and B.S., Investigation: J.L., K.L., and B.S., Visualization: J.L. and B.S., Writing: J.L., K.L, A.V., M.M., P.G., and B.S., Supervision: P.G. and B.S.

## Code availability

The codes for normative analysis and model fitting are available in the GitHub repository: https://github.com/JiaLinLi0822/proj-soft-boundary.

## Data availability

No new data were collected for this study. The macaques data are available from ^34^ upon request.

## Supplementary Information

**Fig S1:**
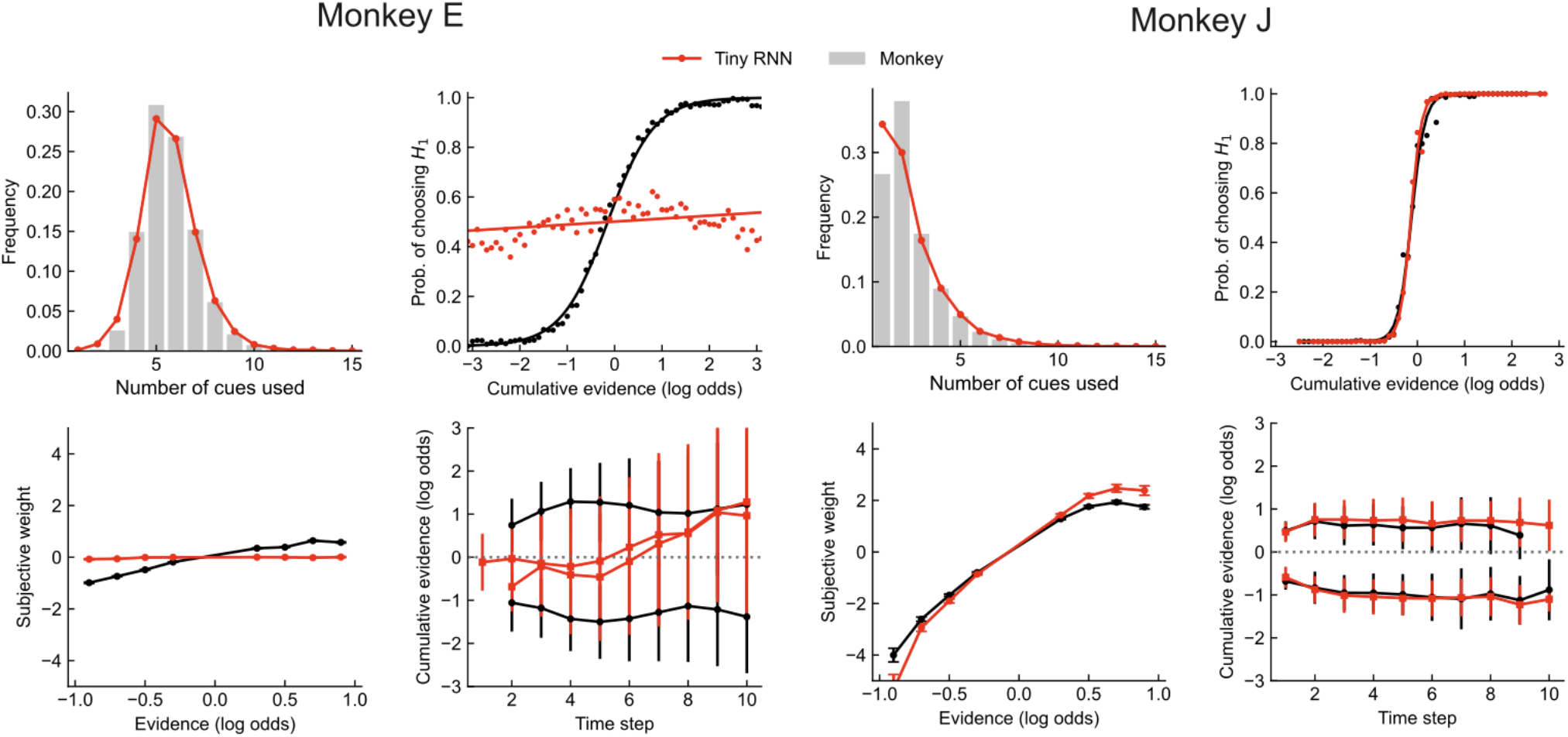
A single-unit tiny RNN fails to fully capture behavioral patterns in non-human primates. Model fits for monkey E (left panels) and monkey J (right panels) using a tiny RNN with one hidden unit. Upper left: distribution of the number of cues used; upper right: psychometric function relating cumulative evidence (logLR) to choice probability; lower left: subjective weights assigned to each stimulus; lower right: cumulative evidence at termination across time.

**Fig S2:**
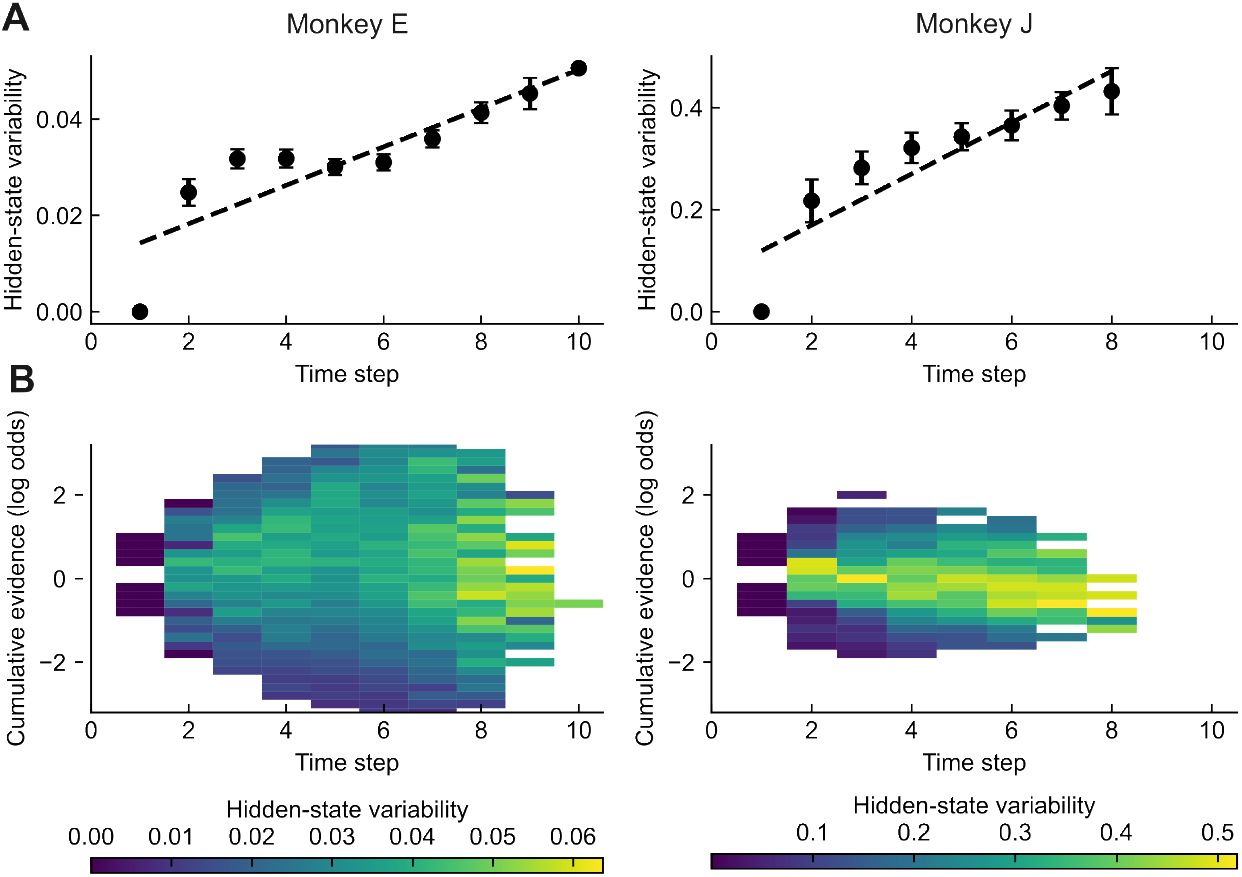
Hidden-state activity variability accumulates over time. (**A**) Variability of the hidden-state activities of the two-unit networks among steps with matched cumulative evidence, as a function of time step, for monkey E (left) and monkey J (right). Dashed line shows the fitted linear regression (Monkey E: β_*t*_ = 3.3*e*^-−3^, *p* < 1*e*^−3^; Monkey J: β_*t*_ = 5.1*e*^−2^, *p* < 1*e*^−3^). (**B**) Variability of hidden-state activities in bins of cumulative evidence over time steps.

## 1 Framing the task as a POMDP

The sequential sampling task in the main text can be framed as a POMDP with two possible world states *w* ∈ {0,1}. Each trial begins with a fixed world state *w* that is unknown to the observer. The observer has to infer the world state from a stream of evidence *e*_*1:t*_:= {*e*_*1*_,…,*e*_*t*_} that is conditionally independent over time given *w* and generated with the state-conditional frequencies or densities ℱ = {*f*_0_, *f*_1_}.

The observer holds hypotheses ℋ *=* {*H*_*0*_, *H*_*1*_} corresponding to the two world states, with an initial belief 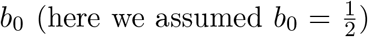. Because the choice is binary, the posterior beliefs of the two hypotheses are complementary,

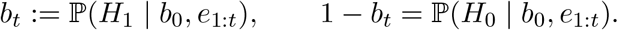

The posterior log-odds between *H*_1_ and *H*_o_ is defined as

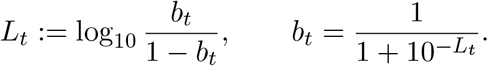

As in the main text, log-odds and log likelihood ratios are expressed in base 10, to align with the literature **? ?**. None of the results below depends on this choice; changing the base only rescales *L*_*t*_, the per-cue log likelihood ratio *l*_*t*_, and the noise scale α by a common constant.

At each time *t*, the observer takes one action *a* from the set *A =* {0,1,*S*}, where 0 means stop and choose *H*_0_, 1 means stop and choose *H*_1_, and *S* means continue sampling at a cost *c* > 0. The observer receives reward *r* > 0 only if they choose the correct world state and receives zero reward otherwise. Therefore, if the observer commits to *a*_*T*_ ∈ {0,1} at time *T* after taking *N*_*s*_ sampling actions, the net reward is

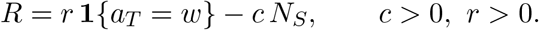

In the task of the main text, the first cue is presented before the first decision, so that *Ns = T —* 1.

## 2 Ideal Bayesian observer shows hard policy boundaries

For an ideal observer with perfect evidence representation, perfect evidence integration, and deterministic action selection, there exist hard decision boundaries 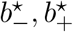 with

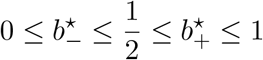

that maximize expected reward with the deterministic threshold form

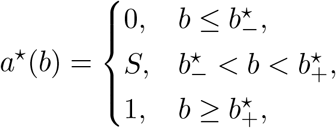

with any fixed deterministic tie-breaking rule at boundary points where two actions have equal value.

Equivalently, in posterior log-odds

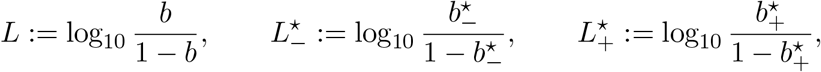

the deterministic threshold form is

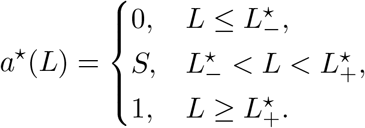

This result is well known in the theory of partially observable Markov decision processes [1, 2]; to better compare with the following analyses under noise, we replicate the proof below to show why hard policy boundaries emerge.

*Proof*. For an ideal observer, ℱ is known to the observer and evidence is integrated perfectly over time without noise. Since the evidence is conditionally independent in time, the belief depends only on the previous belief *b*_t−1_ and the latest piece of evidence *e*_*t*_. Thus, the belief is updated by

Bayes’ rule,

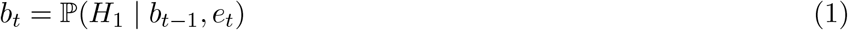

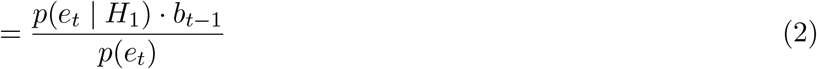

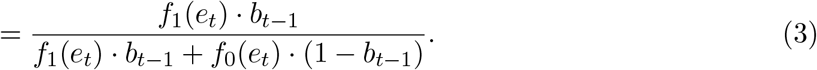

The posterior log-odds is updated as

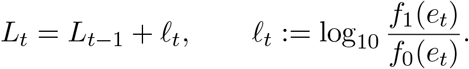

At any stopping time *t*, the conditional expected reward from stopping and choosing *H*_0_ is

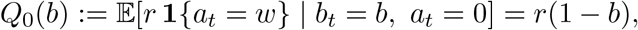

and from stopping and choosing *H*_1_ is

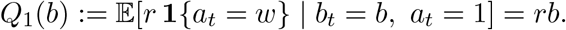

If the agent continues sampling, Bayes’ rule updates the belief to

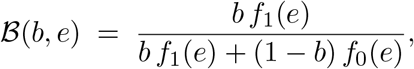

with predictive probability of the evidence

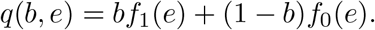

Thus, the expected value of sampling is the predictive-probability-weighted value minus the sampling cost *c*,

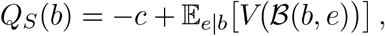

where

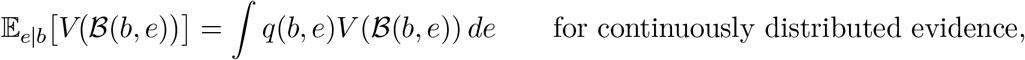

and

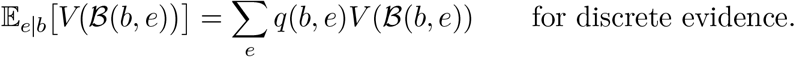

Suppose the observer is allowed to use any mixed action rule π (· | *b)* on the simplex

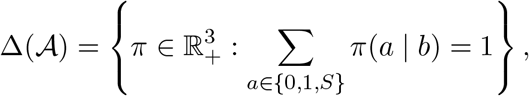

where π*(a* | *b)* denotes the probability of taking action *a* given belief *b*. For a fixed continuation value function *V*, the Bellman backup under the policy π is

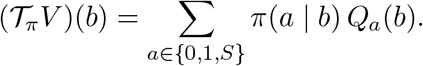

Optimizing this backup gives

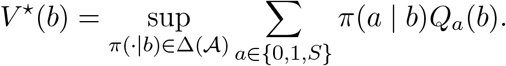

Because the objective is linear in π (·| *b)* and the feasible set is a simplex, the supremum is attained at an extreme point of the simplex, that is, by a Dirac measure on the action with the highest value,

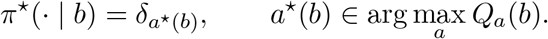

Hence, the optimal value is obtained from a deterministic policy,

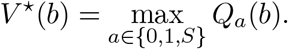

Next, a well-known result from the theory of partially observable Markov decision processes [1, 2] is that *V*\*(*b*) is the limit of a sequence of piecewise linear convex functions *V*_*n*_*(b)* generated by successive application of the Bellman operator. As a result, whenever the limit *V*(b)* exists, it is also a convex function of *b*.

We now use the convexity of *V*(b)* to prove that there exist two deterministic boundaries 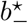 and 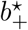 with

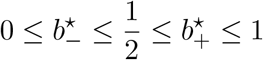

and the threshold form above. First, certainty states are absorbing for an ideal Bayesian observer. If *b* = 0, then *H*_0_ is known with certainty, so stopping immediately yields reward *r*, whereas continuing only adds cost; likewise, if *b* = 1, the agent should stop immediately and choose *H*_1_. Thus

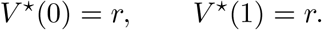

Now, we restrict the belief to the lower half of belief space, 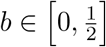 the same conclusion follows for the upper half.

In the lower half, choosing *H*_0_ is always at least as good as choosing *H*_1_. We define *U*_0_ as the difference between the optimal value *V*(b)* and the value of choosing *H*_0_,

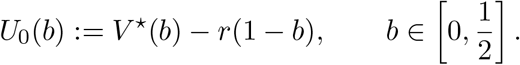

Because *V*(b)* is convex and *r*(1 −*b)* is affine, *U*_0_ is convex on 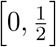. By the definition of the optimal value,

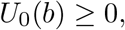

and stopping with action 0 is optimal if and only if

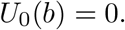

In addition, since *V* ^*\**^(0) = *r*,

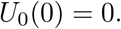

Consider the zero set for stopping,

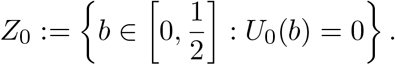

Because *U*_0_ is convex and nonnegative, *Z*_0_ is convex: if *b*_1_, *b*_2_ ∈*Z*_0_ and *α* ∈[0, 1], then

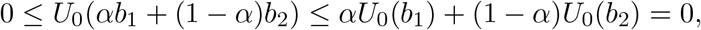

Hence

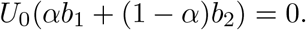

Therefore, all beliefs between *b*_1_ and *b*_2_ also belong to the zero set, and *Z*_0_ is an interval. Since it contains 0, there exists 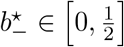 such that

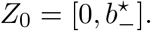

Equivalently,

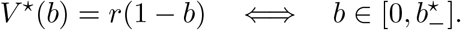

Therefore, throughout 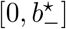, stopping and choosing *H*_0_ is optimal. Moreover, if *U*_0_(*b*_0_) > 0 at some 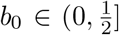, then convexity and *U*_0_(0) = 0 give *U*_0_(*b*) ≥ (*b/b*_0_) *U*_0_(*b*_0_) > 0 for all 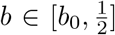.

Thus, once continued sampling is optimal at some belief in the lower half, it remains optimal for all beliefs between that point and 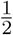.

By the same reasoning for the upper half of belief space, there exists 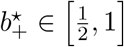 such that

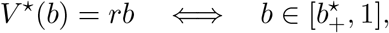

and throughout 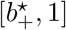, stopping and choosing *H*_1_ is optimal.

For beliefs between these two stopping regions, 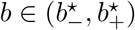, *b* belongs to neither stopping set, and continued sampling attains the optimal value:

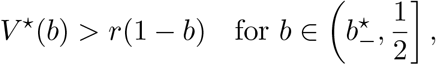

And

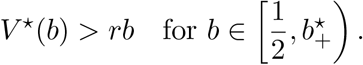

Overall, the optimal policy is

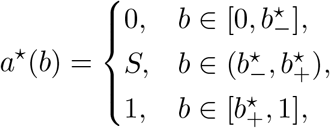

with deterministic tie-breaking permitted at the boundary points. This proves the threshold structure in posterior-probability space.

Because the mapping

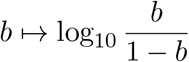

is strictly increasing on (0, 1), the same policy is a threshold rule in log-odds. Let

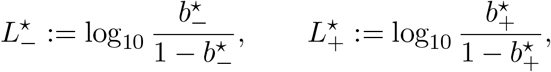

with the conventions 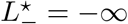 if 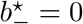 and 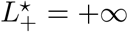 if 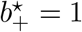.Then

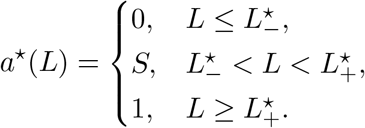

Thus, the ideal observer adopts deterministic decision boundaries.

*Remark (time-constrained condition)*. With a deadline, the value function *V*_*t*_(*b*) is obtained by backward induction from a terminal condition, and each *V*_*t*_ is piecewise linear and convex in *b* [1]. Applying the argument above at each time step yields time-dependent boundaries 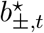, or equivalently 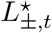, which are hard at every step.

However, biological decision makers are unlikely to meet this idealization, and human and animal behavior is consistently stochastic. The selection of actions could be stochastic rather than deterministic, as described by random utility theory [3–6]. In addition, the brain operates with finite precision in representing each piece of evidence and with imperfect integration of evidence over time. In the following sections, we derive the policies under these constraints, considering separately the effects of stochastic action selection and of noisy or distorted evidence representation.

## 3 Stochastic action selection leads to soft policy boundaries

Suppose tBayesian observer shows hard hat the observer has access to the exact action values

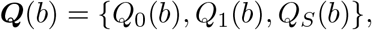

computed from the ground-truth posterior belief *b*, but action selection is stochastic. Specifically, assume that the utility of action *a* ∈{0, 1, *S*} at the moment of selection is

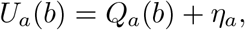

Where

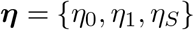

is a random vector with joint distribution *F*_***η***_, and the agent selects the action with the largest *U*_*a*_(*b*).

The resulting policy is generally probabilistic rather than deterministic. For each action *a* ∈ {0, 1, *S*}, the choice probability is

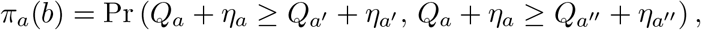

where *a*^*′*^ and *a*^*″*^ denote the two actions other than *a*.

Defining the pairwise perturbation di!erences

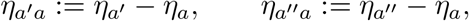

we obtain

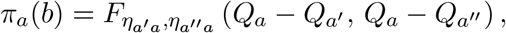

where 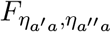 is the joint CDF of the pairwise perturbation differences.

Therefore, even when the underlying value functions retain the deterministic threshold structure of the ideal observer, stochasticity at the stage of action selection transforms the policy into a smooth transition across action boundaries whenever 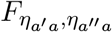 is continuous, rather than a hard threshold rule.

In the special case where the perturbations are i.i.d. type-I extreme-value (Gumbel) variables,

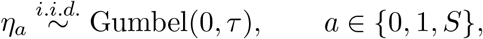

the policy takes the multinomial logit, or softmax, form

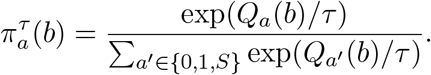

Consequently, the pairwise log-odds between two actions *a* and *a*^*′*^satisfy

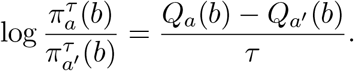

The deterministic indi!erence points, where

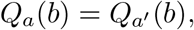

remain the central switching points between pairs of actions, at which 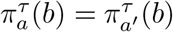. However, the transition around these points is smoothed by the scale *τ*. As *τ*→ 0, the policy converges to the deterministic hard-boundary policy. As *τ* increases, the boundary becomes progressively softer.

Thus, stochastic action selection produces soft decision boundaries even when inference and value computation are otherwise ideal.

*Proof*. Consider the ideal observer, whose action values are computed from the ground-truth belief,

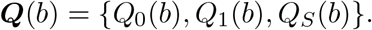

A source of stochasticity at the moment of action selection perturbs these values,

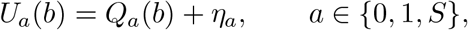

Where

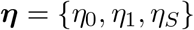

is a random vector with joint distribution *F*_***η***_.

For a commonly assumed case for categorical choices, let the perturbations be i.i.d. type-I extremevalue (Gumbel) variables [4],

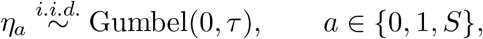

with CDF

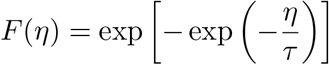

and density

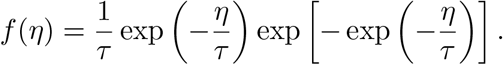

Using the independence of the three perturbations,

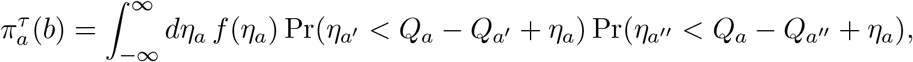

that is,

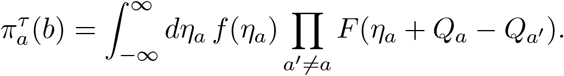

Substituting the Gumbel density and CDF gives

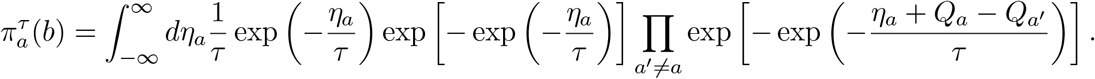

Since

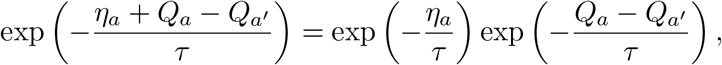

we obtain

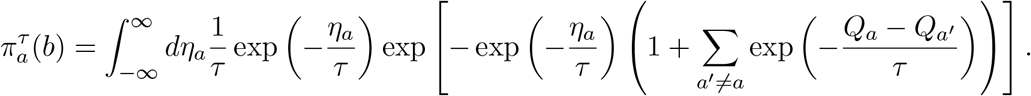

Let

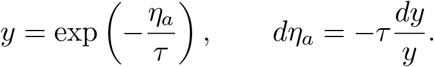

As *η*_*a*_→− ∞,*y*→∞ and as *η*_*a*_→ ∞

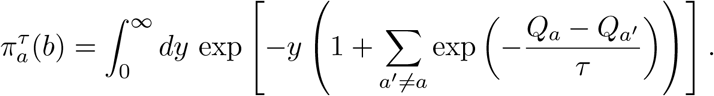

Using 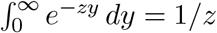 for *z*>0, we get

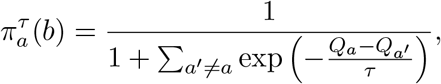

and, rearranging,

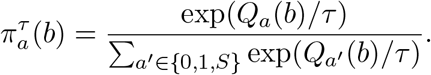

This is the multinomial logit, or softmax, policy. The pairwise log-odds are

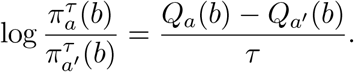

Thus, the deterministic indifference points 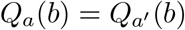 remain the central switching points, but the transition around them is smooth rather than discontinuous. This concludes the proof that stochastic action selection produces soft decision boundaries.

We note that the connection between Gumbel perturbations and softmax actions has been described and used extensively in the machine learning literature [7].

## 4 Soft policy is optimal under costly control

In the previous section, we assumed stochastic action selection and showed that, even if the agent knows the true action values ***Q***(*b*), it behaves with soft policy boundaries,

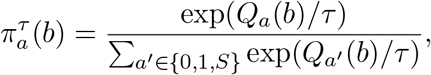

where *τ* is the scale parameter of the Gumbel perturbations.

In this section, we show that the cost of control, as motivated by information theory and thermo-dynamics [8–10], provides a normative basis for this stochastic action selection. Specifically, we use the Kullback–Leibler (KL) divergence of the policy from a uniformly random default policy as the cost of control, and we subtract this cost from the expected gain to identify the optimal policy under costly control.

We prove that the optimal policy has soft boundaries without any assumption about the distribution of perturbations in action selection. In other words, if the agent maximizes expected gain net of the cost of control, it behaves as if its action values were perturbed by Gumbel noise.

Assume that an agent chooses among actions *a* ∈*A*, where *A* is a finite action set with *N* elements (*A* = {0, 1, *S*} and *N* = 3 in the main text). Assume further that the agent knows the true action values *Q*_*a*_(*b*) at its current belief *b*. The agent chooses a policy π_*a*_(*b*) *≥* 0 that assigns a probability to each action in *A*, such that

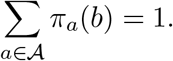

The expected gain of this policy is

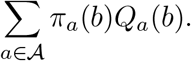

We now assume a cost associated with implementing a more precise policy. To describe this cost, we first choose a default policy that the agent adjusts through control. sThe natural choice is the uniformly random policy over all actions, 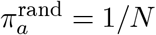.

Following the thermodynamics literature [8–10], the cost of control of a policy ***π*** is defined as its KL divergence from the default policy,

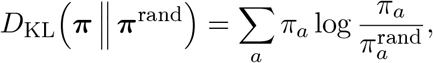

measured in nats. For the uniform default policy, this quantity equals the reduction in entropy relative to the random policy, *D*_KL_ = log *N− H*(***π***). Importantly, this cost does not depend on the mechanism by which control is exerted; it arises from the entropic cost of reducing uncertainty in the policy. The overall value of a policy, taking this cost into account, is

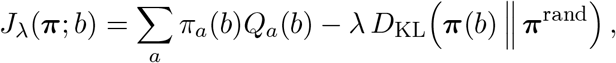

where the constant *λ* > 0, the price of control, converts information measured in nats into units of reward and measures how efficiently the agent can exert control over its policy. Thus, the agent chooses π_*a*_(*b*) to maximize

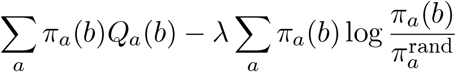

subject to the constraint ∑_*a*_ *π*_*a*_(*b*) = 1. We find the maximum with a Lagrange multiplier *µ* for the constraint,

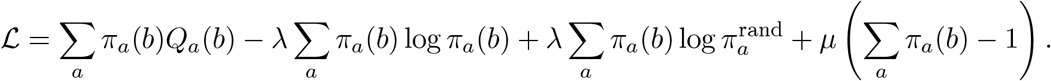

Taking the derivative with respect to *π*_*a*_(*b*) and setting it to zero gives

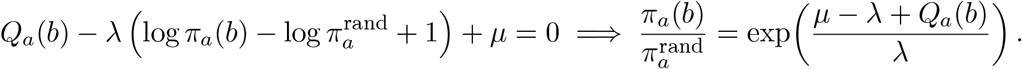

Since *µ* is a constant independent of *a* and 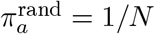, we set 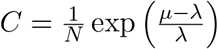 and write

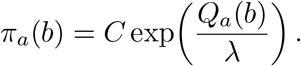

The normalization condition determines *C* and yields

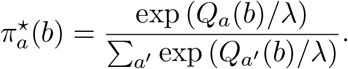

Because *J*_*α*_ is strictly concave in ***π*** (a linear term minus *λ* times a strictly convex KL term), this stationary point is the unique maximum. It is exactly the softmax policy obtained from Gumbel perturbations, with the scale of the perturbations equal to the price of control, *τ* = *λ*. Thus, the cost of control leads to soft policy boundaries exactly as stochastic action selection does.

This proof is agnostic about the mechanism by which the agent trades o! the cost of control against the higher expected gain of a more precise policy. Whenever there is such a cost, the agent should adopt soft boundaries, with softness set by the price of control *λ*.

## 5 Belief noise makes hard policy boundaries appear soft

To model non-ideal inference, let the observer represent each piece of evidence with finite precision and integrate evidence imperfectly over time. A general formulation of the update of the internal log-odds and the corresponding internal belief is

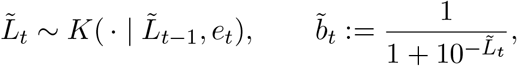

where the kernel *K* encodes noisy representation, misweighting of cues, quantization, leak, saturation, or other losses during integration. There exist hard decision boundaries 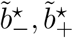 in the observer’s internal belief with

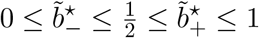

that are optimal, with the deterministic threshold form

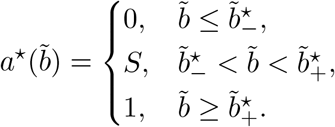

Equivalently, in log-odds coordinates,

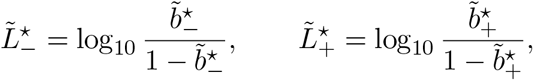

so that

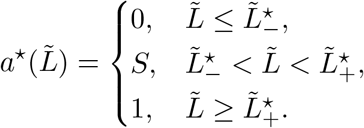

In the time-constrained condition, the boundaries become time dependent, 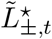.

The experimenter, however, observes soft decision boundaries when plotting actions against the ground-truth log-odds *L*_*t*_:

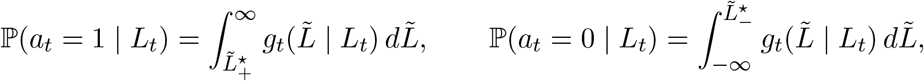

and

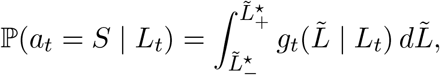

where *g*_*t*_ is the conditional law of the observer’s internal log-odds given the ground-truth log-odds,

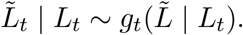

In the scalar case with Gaussian noise,

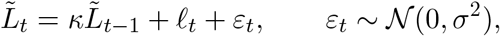

where *k∈* [0,1] is the retention of the previous internal state *(k* = 1 for perfect integration and *K* < 1 for a leak), *l*_*t*_ is the ground-truth log likelihood ratio of evidence *e*_*t*_, and *ε*_*t*_ is i.i.d. Gaussian representation noise, the internal log-odds given the evidence sequence is distributed as

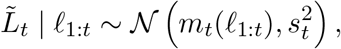

Where

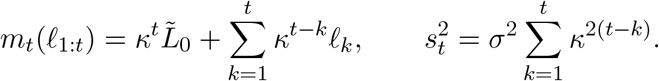

The experimenter’s observed choice probabilities are

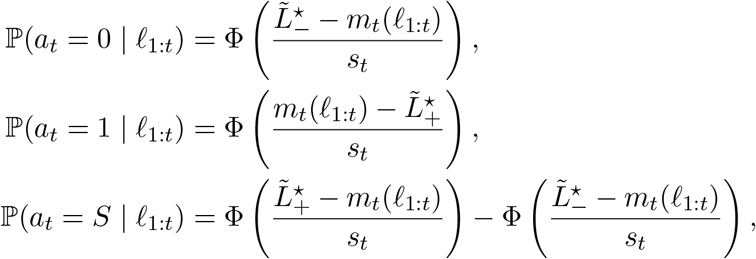

which show soft boundaries in the experimenter’s observed choice probabilities, with softness set by the dispersion *s*_*t*_ and hence by the noise scale σ and the retention *k*. For perfect integration *(k =* 1) from an unbiased start 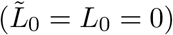, *m*_*t*_ *= L*_*t*_ and 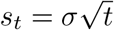; this is the belief-noise model of the main text, 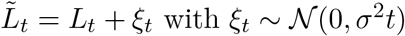.

*Proof*. Write the general update of the observer’s internal log-odds as

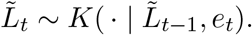

The internal belief of the observer is

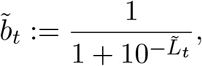

and the internal action values of the three actions *a* ∈ {0,1, *S*} are

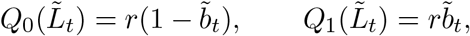

and

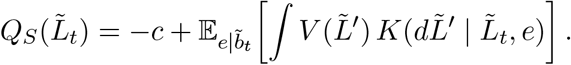

Specifically,

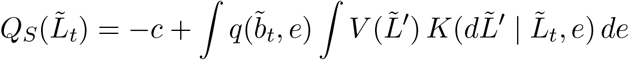

for continuously distributed evidence, and

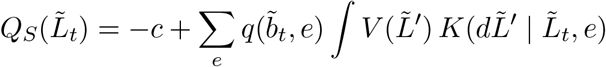

for discrete evidence, where

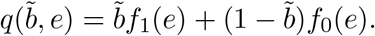

The rest of the proof follows the ideal-observer case. Suppose the agent may use any mixed action rule 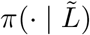 on the simplex

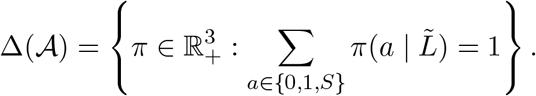

For a fixed continuation value function *V*, the Bellman backup at state 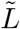 is

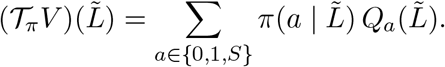

Therefore the optimal Bellman backup is

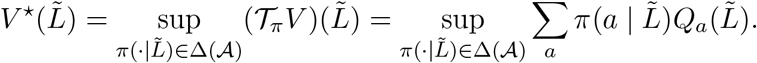

Because the objective is linear in 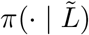 and the feasible set is a simplex, the supremum is attained at an extreme point of the simplex. Hence

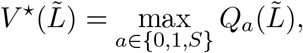

and any optimal policy is deterministic in the observer’s internal state,

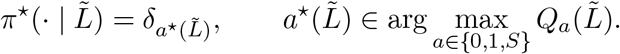

Following the same arguments as for the ideal observer, there exist thresholds

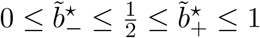

such that

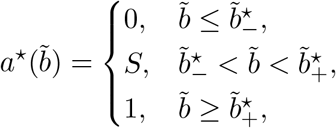

or, in log-odds coordinates,

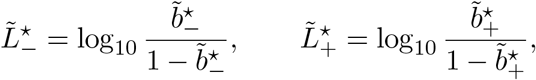

so that

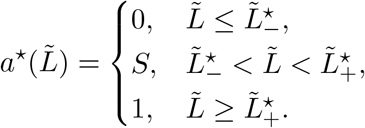

The critical point is that the optimal policy still has hard boundaries in the observer’s internal belief. The experimenter, however, observes soft decision boundaries when plotting actions against

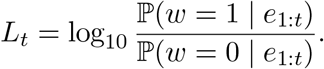

Let the conditional law of the observer’s internal log-odds given the ground-truth log-odds be

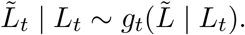

If the internal policy is the hard rule with thresholds 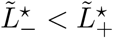, then the experimenter observes

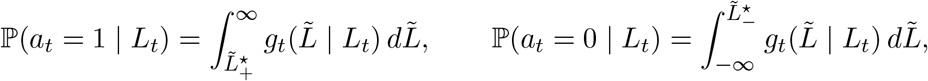

and

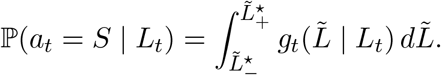

These are smooth functions of *L*_*t*_ whenever *g*_*t*_(· | *L*_*t*_) has a smooth density. Thus, a hard boundary in internal space is generically observed as a soft boundary on the ground-truth evidence axis.

In the scalar case with Gaussian noise,

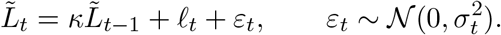

Iterating this recursion gives

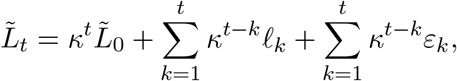

where 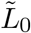 is the initial internal state of the observer.

If the noise terms are independent over time, then, conditional on the evidence sequence,

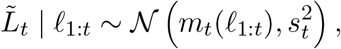

Where

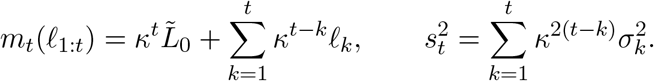

For constant noise variance 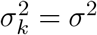,

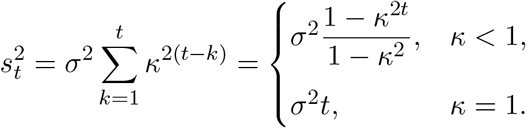

The experimenter’s observed choice probabilities are then

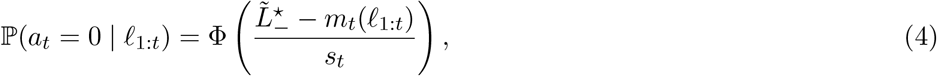

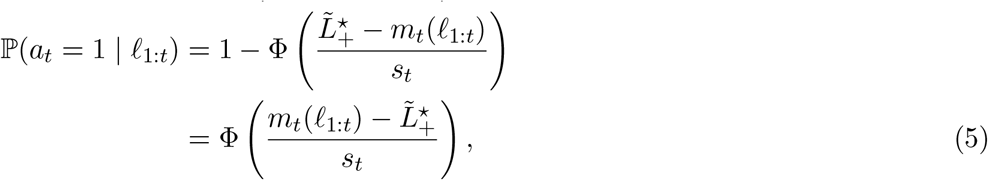

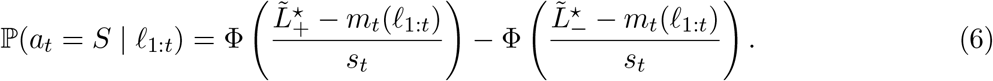

For perfect integration *(k =* 1) from an unbiased start 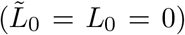, 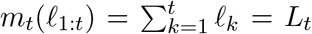 and 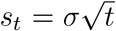, so the internal log-odds is 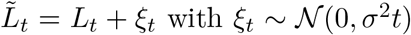, the belief-noise model of the main text. The dispersion *s*_*t*_ of the internal evidence around the ground truth then grows with every sample, and the observed policy broadens over time. As this form of the solution shows, the experimenter observes soft boundaries despite the observer’s hard-boundary strategy in its own belief space.

*Remark (distorted accumulation as belief noise)*. The kernel *K* need not be stochastic. Suppose the observer integrates evidence without noise but with subjective weights 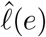 that differ from the true log likelihood ratios 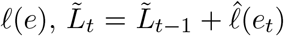. Conditional on the full evidence sequence, 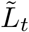 is then deterministic. The experimenter, however, conditions on the ground-truth log-odds 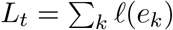, and different evidence sequences with the same *L*_*t*_ yield different 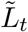 whenever 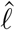 is not proportional to *l*. The conditional law *g*_*t*_(· | *L*_*t*_) is therefore non-degenerate, and the observed policy is soft by the argument above. Because more sequences with different internal values map onto each value of *L*_*t*_ as evidence accumulates, this dispersion generally grows with *t*. From the experimenter’s perspective, systematic misweighting of cues and random representation noise thus both constitute belief noise: both corrupt the belief upstream of the policy, and both produce a dispersion of the internal belief at fixed ground-truth evidence that accumulates over samples.

